# Detection of incipient pancreatic cancer with novel tumor-specific antibodies in mouse models

**DOI:** 10.1101/2020.09.10.292193

**Authors:** Tobiloba E. Oni, Carmelita Bautista, Joseph R. Merrill, Jeroen A.C.M. Goos, Keith D. Rivera, Koji Miyabayashi, Giulia Biffi, Libia Garcia, Dennis Plenker, Hardik Patel, Ela Elyada, Maria Samaritano, Kenneth H. Yu, Darryl J. Pappin, Michael G. Goggins, Ralph H. Hruban, Jason S. Lewis, Scott K. Lyons, Johannes T. Yeh, David A. Tuveson

**Author notes:** Correspondence to David A. Tuveson: 1 Bungtown Rd, Cold Spring Harbor, NY 11724.

## Abstract

Pancreatic ductal adenocarcinoma (PDAC) is a highly lethal malignancy, as 90% of patients do not survive beyond five years from diagnosis. This dismal prognosis is largely due to the advanced stage of the disease at diagnosis, which precludes potentially curative surgical resection. Although early detection strategies hold significant promise for improving patient outcomes, there is still no accurate diagnostic tool to detect incipient PDAC. Here, we sought to develop antibodies for the early detection of PDAC by positron-emission tomography (PET) imaging. Accordingly, we establish a pipeline to generate novel tumor-specific monoclonal antibodies (mAbs) against cell-surface proteins of PDAC patient-derived organoids (PDOs). We identify a panel of 16 tumor organoid-binding antibodies (TOBi-bodies) that display high reactivity to human PDAC tissues but not to matched adjacent normal pancreas. We then employ biochemical, flow cytometric, mass spectrometric, and CRISPR/Cas9-mediated knockout methods to determine the cognate antigens of these TOBi-bodies. We identify two mAbs that bind to tumor-specific variants of the surface protein CEACAM6 and show minimal binding to normal tissues. PET imaging in mouse models using these TOBi-bodies enables the detection of incipient human organoid-derived PDAC tumors that are rather undetectable by palpation or high-resolution ultrasound imaging techniques. We propose that further development of these mAbs as PET radiotracers could facilitate the early detection and accurate staging of PDAC.

## Introduction

Pancreatic ductal adenocarcinoma (PDAC) is the most lethal common malignancy, with a 5-year survival rate of less than 10% [1]. This abysmal prognosis reflects the advanced stage of the disease at diagnosis. Indeed, due to the asymptomatic nature of incipient PDAC, most patients (80-85%) present with metastatic or locally advanced disease, which precludes surgical resection and leaves relatively ineffective chemotherapies as the only treatment option [2]. For the minority of patients diagnosed with early stage, localized disease, surgery combined with chemotherapy prolongs survival and offers the best chance for a cure [2]. Therefore, detecting PDAC at an early stage would increase the proportion of surgical candidates and significantly improve outcomes in this deadly disease [2]. Despite the potential impact of early detection on patient outcomes, there are currently no clinical tools to non-invasively detect early stage PDAC. Consequently, effective early detection methods remain one of the most important unmet needs in PDAC.

Imaging plays a decisive role in the diagnosis and staging of PDAC [3]. However, current imaging methods have significant limitations in identifying early disease [4]. Computed tomography (CT) is the most widely used imaging tool for PDAC, but it lacks sensitivity to detect small tumors (<2cm) and micro-metastases [5]. Similarly, magnetic resonance imaging (MRI) displays low accuracy for identifying small tumors [6]. Although endoscopic ultrasound (EUS) exhibits high sensitivity for detecting lesions between 1-2 cm [4], EUS is invasive, suffers from operator-dependent variability in image acquisition and interpretation, and has a limited role in the detection of occult metastasis [4]. In contrast to EUS, positron emission tomography (PET) using the glucose analog, 18F-fluorodeoxyglucose (FDG), enables wide anatomical coverage and simultaneous detection of primary and metastatic disease [7]. However, FDG uptake is not tumor-specific since inflammatory diseases, such as chronic pancreatitis, exhibit PET-avidity comparable to tumors [8].

Immuno-positron emission tomography (immuno-PET) combines the exquisite specificity of antibodies with the inherent sensitivity of PET imaging via the conjugation of antibodies to radionuclides [9]. The integration of CT with immuno-PET imaging (immuno-PET/CT) rectifies the low spatial resolution of PET imaging and enables the delineation of proximal anatomical structures [10]. This dual modality is under active investigation in PDAC. A recent phase 1 clinical study compared conventional CT imaging to immuno-PET/CT using a zirconium-labeled antibody against the tumor marker, CA19-9 in PDAC patients [11]. Remarkably, immuno-PET/CT identified small metastatic lesions (<1 cm) that were not detectable by conventional CT imaging, indicating that immuno-PET imaging can delineate occult metastasis and improve the accuracy of preoperative staging. However, as CA19-9 is also highly expressed in benign diseases [12], radiolabeled antibodies targeting this antigen will yield false positives. More so, since 5-10% of Caucasians and 22% of African Americans and Africans cannot synthesize CA19-9 [12], CA19-9-based radiotracers will yield false negatives in these individuals and are thereby unsuitable for the diagnosis of PDAC. Therefore, to accurately detect incipient PDAC and occult metastatic disease, tumor-specific antibodies must be developed for immuno-PET imaging. These tumor-specific radiotracers will enable early PDAC detection, thereby facilitating timely surgical intervention and prolonging patient survival.

Since the development of the hybridoma method for producing monoclonal antibodies (mAbs), numerous studies have focused on developing tumor-specific antibodies to PDAC [13-15]. However, none of these studies have identified mAbs with exclusive binding to malignant cells. Since these studies mostly used monolayer PDAC cell lines as immunogens, and cell lines poorly recapitulate the inter- and intra-tumoral heterogeneity found in patients [16], a decrease in the expression of shared PDAC antigens in cell lines could potentially explain previous unsuccessful attempts.

Here we sought to develop tumor-specific antibodies for the detection of early PDAC by immuno-PET imaging. To this end, we produced rat hybridomas against membrane proteins extracted from a pool of patient-derived organoids (PDOs). As PDOs retain the heterogeneity of the original tumors [17], these ex-vivo cultures contain a more diverse antigen representation than cell lines and thereby more accurately reflect the set of shared PDAC antigens. Using flow cytometry, we first identified tumor-organoid binding antibodies (TOBi-bodies) that exhibited binding to accessible antigens on the tumor cell surface. Using immunohistochemistry (IHC) on human PDAC and matched adjacent normal pancreas tissues, we then identified 16 tumor-selective TOBi-bodies. Importantly, we discovered two TOBi-bodies that displayed strong binding to PDAC but showed minimal binding to chronic pancreatitis samples and various normal tissues. Immuno-PET imaging using these TOBi-bodies enabled the detection of incipient human organoid-derived PDAC tumors in orthotopic mouse models. Our data indicate that these TOBi-bodies could be developed as non-invasive PET radiotracers for early PDAC detection.

## Results

### Generation of Tumor-selective Antibodies

To develop tumor-specific antibodies to PDAC, we established a pipeline with the goal of identifying mAbs that bind cell-surface antigens in human PDAC tumors but not in matched normal pancreas (Fig. 1A). First, we cultured organoids derived from 15 patients with PDAC [17] (Table S1), harvested and then pooled the samples. Next, we extracted membrane fractions from the pooled samples and immunized rats with this extract. Following four injections in two-week intervals, we collected blood from the immunized rats and assayed antibody serum titers to the human PDAC SUIT2 cell line by cell-based Enzyme-linked Immunosorbent Assay (ELISA) (Fig. S1A). Although we immunized the rats with human organoids, cell lines are more amenable to cell-based ELISA strategies. We also expected some degree of overlap in the cell-surface antigen representation between PDAC cell lines and organoids. Particularly, we chose SUIT2 because it contains frequently altered mutations in PDAC (*KRAS*^G12D^; *TP53*^R273H^; *CDKN2A*^E69*^) and possesses a hybrid molecular subtype, with the expression of both classical and basal-like genes [18]. Therefore, titering the sera of immunized rats on SUIT2 cells should approximate the antibody response to the panel of PDOs. As expected, serum IgGs from rats immunized with PDO membrane extracts displayed concentration-dependent binding to SUIT2 cells by cell-based ELISA, whereas control sera from phosphate-buffered saline (PBS)-immunized rats displayed no binding (Fig. S1B). These results demonstrate that membrane extracts from PDOs elicited robust humoral response and that the antibody repertoire of PDO-immunized rats contains mAbs against human PDAC.

**Figure 1.**
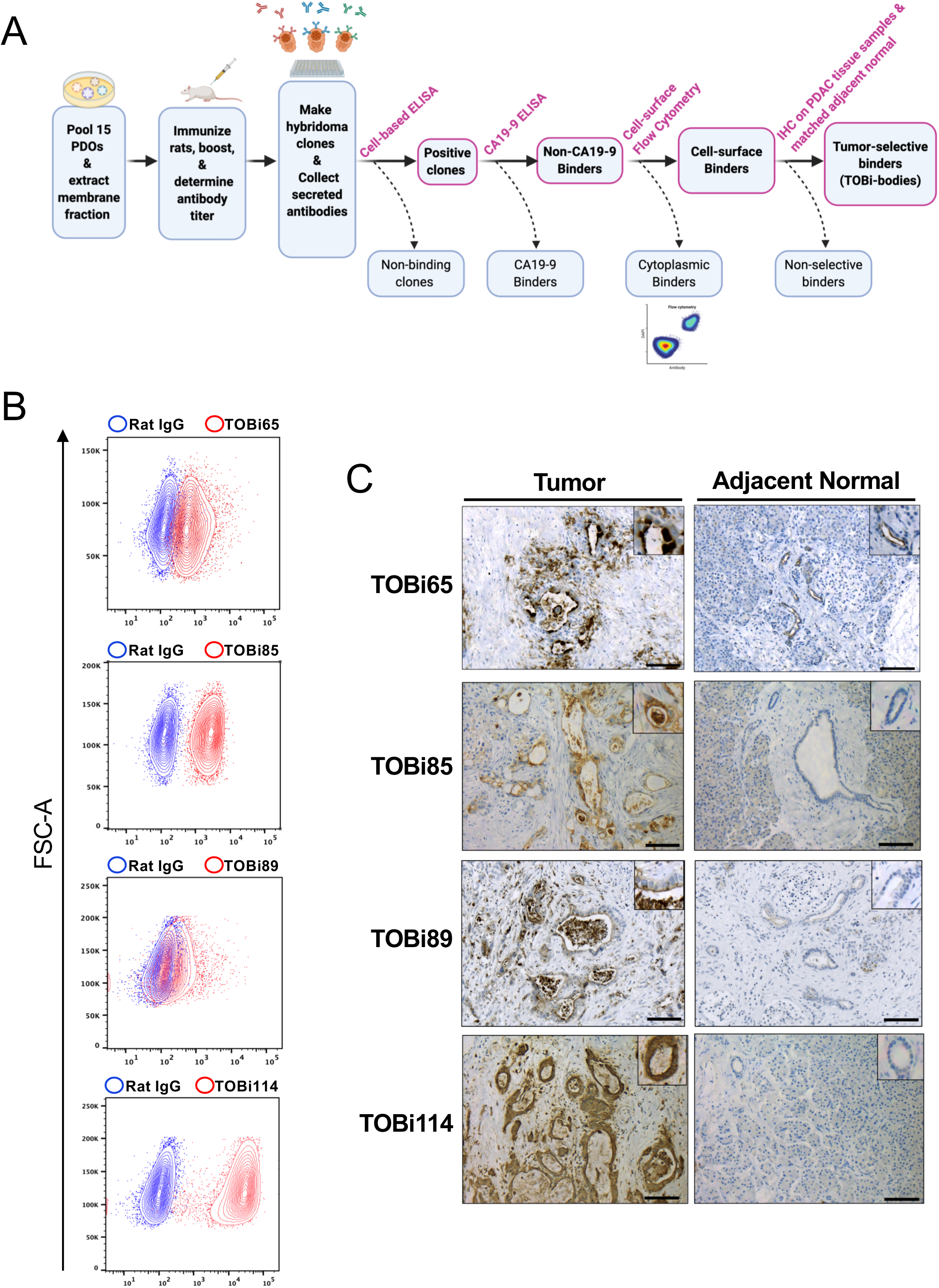
Generation of tumor-selective antibodies. (*A*) Screening pipeline for generating tumor-selective antibodies in rats (n=3). PDOs, patient-derived organoids; ELISA, enzyme-linked immunosorbent assay; IHC, immunohistochemistry; TOBi-bodies, tumor-organoid binding antibodies. (*B*) Candidate antibodies (red) display cell-surface binding compared to rat IgG control (blue) in SUIT2 cells by cell-surface flow cytometric analysis. Results are representative of three independent experiments. FSC-A, forward scatter-area. (*C*) Immunohistochemical labeling of human PDAC (left) and adjacent normal tissue sections (right) with TOBi-bodies. Images are representative of three biological replicates. Inserts: magnification. Scale bars, 100 μm.

We next extracted splenocytes from the immunized rats and performed fusion with rat myeloma cells to generate hybridomas. We established 1356 rat hybridomas fusion clones and collected hybridoma conditioned media containing secreted mAbs. To identify hybridoma clones secreting tumor-reactive mAbs, we tested the reactivity of hybridoma supernatant on SUIT2 cells by cell-based ELISA. By comparing the binding of mAb-containing hybridoma supernatants to rat IgG antibody controls, we identified several clones with increased binding to SUIT2 cells (Fig. S1C). To select positive clones, we used a low cut-off of 1.5-fold compared to control IgG antibodies in order to include mAbs that bind antigens that may be highly expressed in PDOs but show reduced expression in SUIT2 cells. Based on this cut-off, we found that 251 out of 1356 clones (18.5%) produced SUIT2-reactive mAbs (Fig. S1C). Subsequently, we expanded these positive clones to produce larger quantities of mAbs; however, only 77 out of the 251 hybridomas remained viable after passaging. This high attrition is common and has been attributed to chromosomal loss induced by the fusion process [19]. Since CA19-9 is a well-characterized PDAC antigen with poor tumor-specificity, we next sought to exclude CA19-9-reactive mAbs. We performed CA19-9 ELISA with hybridoma supernatants from all 77 clones and found that 12 clones secreted mAbs reactive to CA19-9 (Fig. S1D) and 65 mAbs were negative for CA19-9 binding.

In line with our aim of generating antibodies to cell surface-accessible tumor epitopes, we next sought to determine whether the 65 candidate mAbs reacted with cell surface antigens. To this end we stained SUIT2 cells with negative control rat IgG or candidate mAbs, and 4’,6-diamidino-2-phenylindole (DAPI) to enable the exclusion of non-viable cells. We then performed flow cytometric analysis to identify clones that exhibited elevated extracellular binding compared to rat IgG negative control. Accordingly, we found that 49 out of all 65 mAbs were cell-surface binders with varying binding intensity (Fig. 1B, S2 and S3). The variation in binding intensity may reflect the binding of antigens or epitopes with different levels of expression, or it may be due to differences in antibody affinity.

These 49 mAbs display cell-surface binding to PDAC cells *in vitro*. However, since the diagnostic utility of antibodies depends on their ability to distinguish normal cells from tumor cells, we sought to determine whether the candidate mAbs were tumor-selective. To this end, we performed immunohistochemistry (IHC) to compare the reactivity of these mAbs to PDAC tumors and matched adjacent normal pancreatic tissues. Rat IgG control antibodies displayed no labeling in these tissues (Fig. S4). Therefore, positive labeling observed with these mAbs should reflect antigen specificity. Using this approach, we identified 16 mAbs, subsequently referred to as tumor organoid binding antibodies (TOBi-bodies), with elevated binding in neoplastic cells, but low or no binding in matched adjacent normal samples (Fig. 1C and S4).

Overall, we established 1356 hybridomas. 251 clones were positive for binding in our cell-based ELISA. 77 hybridomas survived passaging and expansion, and we deprioritized 12 CA19-9 binders. We tested the 65 non-CA19-9 clones for cell surface binding in SUIT2 cells and confirmed 49 cell surface binders. Of these antibody clones, 16 mAbs (TOBi-bodies) were specific for PDAC tissue samples. This result validates our approach and indicates that the development of antibodies to PDAC PDOs can generate tumor-selective binders with potential utility for diagnostic applications.

### Identification of the Cognate Antigens of the TOBi-bodies

To determine the cognate antigens of these TOBi-bodies, we first focused on 3 TOBi-bodies with similar flow cytometric binding profiles— TOBi109, TOBi111, and TOBi125 (Fig. S2). To this end, we performed immunoprecipitation in SUIT2 cells using rat IgG control antibody, TOBi109, TOBi111, or TOBi125, followed by isobaric tags for relative and absolute quantification (iTRAQ)-based mass spectrometry (IP/MS). We then ranked the most enriched proteins relative to IgG control and prioritized known membrane proteins. Interestingly, all three TOBi-bodies enriched for CEACAM6 and closely-related CEACAM5, indicating that these antibodies may bind to both glycoproteins (Table S2). Similar analysis with TOBi85 in BXPC3 and MIAPACA2 PDAC cell lines and with TOBi89 in hM19D PDOs revealed CEACAM6 as the most enriched membrane protein and the likely cognate antigen of both TOBi-bodies (Fig. 2A and 2B, Table S3 and S4).

**Figure 2.**
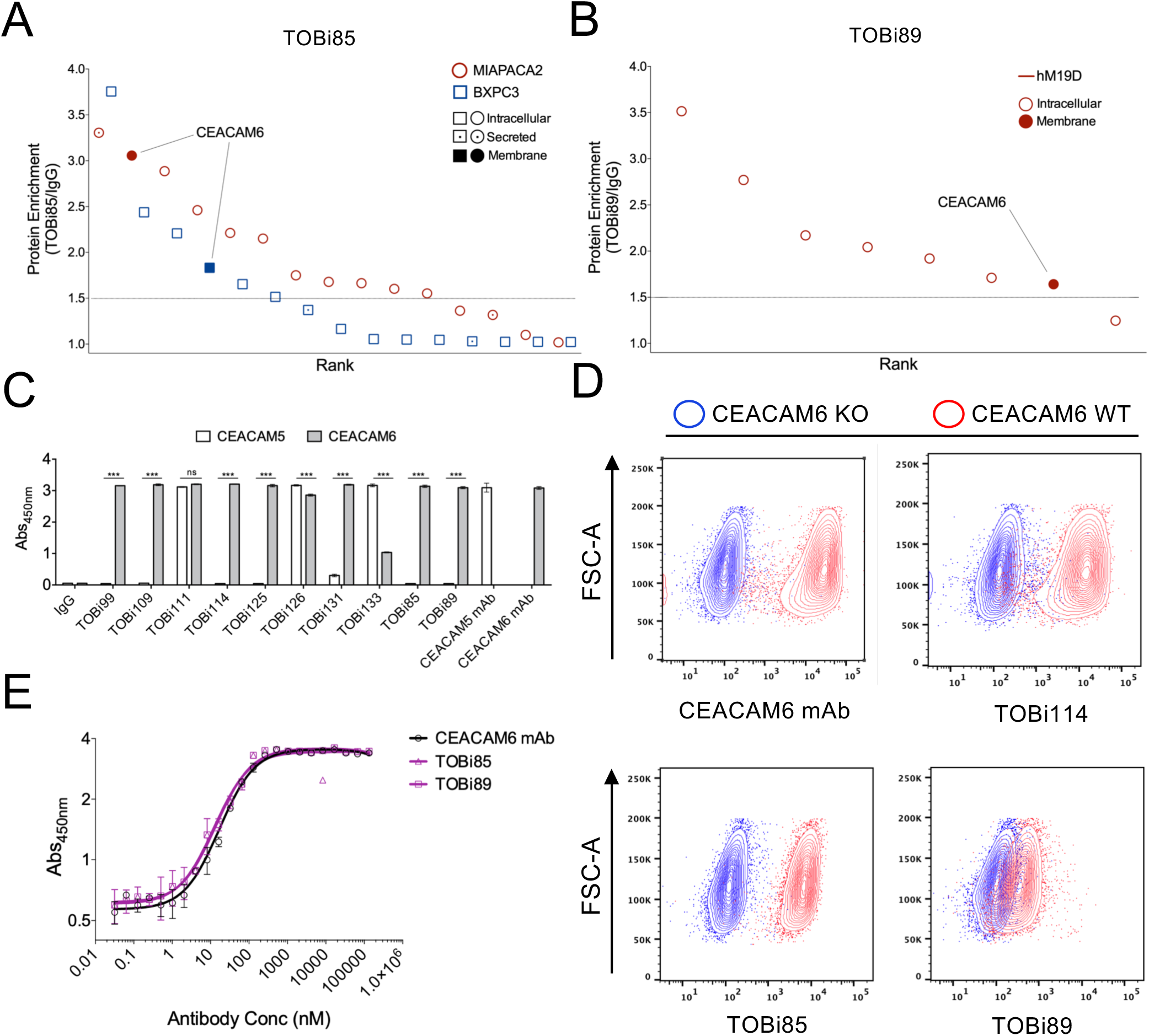
CEACAM6 is the cognate antigen of various TOBi-bodies. (*A*) TOBi85 binds to CEACAM6. Isobaric tags for relative and absolute quantitation (iTRAQ) mass spectrometry was used to determine the abundance of proteins immunoprecipitated by TOBi85 or rat IgG in BXPC3 and MIAPACA2 cells. Enrichment was determined by calculating ratios of protein abundance in TOBi85 relative to IgG-precipitated samples and ranked from the highest to lowest enriched proteins. An enrichment cutoff of 1.5 was applied (dotted line). Subcellular location of enriched proteins was denoted based on UNIPROT annotations, and membrane proteins were prioritized. CEACAM6 is the only membrane protein enriched in both MIAPACA2 and BXPC3. (*B*) TOBi89 also binds to CEACAM6. Immunoprecipitation by TOBi89 or rat IgG in hM19D organoids followed by similar analyses described for TOBi85 (*A*) reveals CEACAM6 as the only enriched membrane protein. (*C*) Various TOBi-bodies bind to CEACAM6 or to both CEACAM6 and CEACAM5. Solid-phase ELISA measurements (absorbance at 450 nm) of the binding of various TOBi-bodies to recombinant CEACAM6 (gray bars) and CEACAM5 (white bars). As expected, rat IgG control displayed no binding, whereas commercially available CEACAM6 and CEACAM5 antibodies displayed specific binding to their antigens. Data are shown as mean ± SD of triplicate wells and are representative of three independent experiments; ***p<0.001, ns (not significant). (*D*) CRISPR/Cas9 knockout of CEACAM6 in SUIT2 cells abolishes the binding of TOBi85, TOBi89, and TOBi114. Cell-surface flow cytometry analysis of CEACAM6 knockout (KO, blue) and wildtype (WT, red) SUIT2 cells using a commercially available CEACAM6 antibody (CEACAM6 mAb, top left), TOBi114 (top right), TOBi85 (bottom left), and TOBi89 (bottom right). Results are representative of three independent experiments. FSC-A, forward scatter-area. (*E*) The unique binding profiles of TOBi85 and TOBi89 are not due to differences in binding affinity to CEACAM6. Serial 10-fold antibody dilution followed by solid-phase ELISA using recombinant CEACAM6 was used to generate affinity profiles. Comparison of the affinity profiles of TOBi85 and TOBi89 (purple) to a CEACAM6 antibody (black) revealed no differences in CEACAM6 binding. Data are shown as mean ± SD of triplicate wells and are representative of three independent experiments.

Next, we sought to validate these findings by determining whether these TOBi-bodies directly bind to recombinantly expressed CEACAM6 and CEACAM5 proteins. Additionally, since the TOBi-bodies may share antigenic determinants, we asked whether other TOBi-bodies also bind to CEACAM6 and CEACAM5. We therefore tested the reactivity of these mAbs with recombinant CEACAM6 and CEACAM5 using solid-phase ELISA and compared their binding with rat IgG and commercially available CEACAM5 and CEACAM6 antibodies. As expected, rat IgG showed no reactivity to CEACAM5 and CEACAM6, whereas positive control antibodies against these proteins showed specific binding to their respective antigens. Interestingly, we found that several TOBi-bodies exclusively displayed binding to CEACAM6 (TOBi85, TOBi89, TOBi99, TOBi109, TOBi114, and TOBi125) or both CEACAM6 and CEACAM5 with a variable degree of reactivity (TOBi111, TOBi126, TOBi131, and TOBi133) (Fig. 2C). Given that mAbs bind to only one epitope, the differences in the cross-reactivity to CEACAM5 among these TOBi-bodies may reflect differences in their cognate epitopes.

TOBi85 and TOBi89 have distinct flow cytometric binding profiles compared to other CEACAM6-binding TOBi-bodies (Fig. 1B and S2). In particular, TOBi89 showed a significantly lower binding intensity to SUIT2 cells. To eliminate the possibility that these mAbs bind to a non-CEACAM6 antigen and merely cross-react with CEACAM6, we performed CRISPR/Cas9 knockout of *CEACAM6* in SUIT2 cells and then evaluated the binding of a commercially available CEACAM6 antibody (CEACAM6 mAb) and CEACAM6-reactive TOBi-bodies, TOBi85, TOBi89, and TOBi114. As expected, *CEACAM6* knockout abrogated the binding of CEACAM6 mAb, validating the loss of CEACAM6 (Fig. 2D). Notably, knockout of *CEACAM6* also abolished the binding of all three CEACAM6-reactive TOBi-bodies (Fig. 2D), corroborating the IP/MS and ELISA data (Fig. 2A, 2B, and 2C). These results collectively confirm that TOBi85 and TOBi89 are CEACAM6 binders despite their distinct flow cytometric binding profiles.

Another factor that may explain the unique binding profiles of TOBi85 and TOBi89 is differences in binding affinity to CEACAM6. To test whether these mAbs have lower binding affinities to CEACAM6, we compared the binding affinities of CEACAM6 mAb, TOBi85 and TOBi89 by ELISA using recombinant CEACAM6. Surprisingly, there were no significant differences in the binding affinities of these mAbs to recombinant CEACAM6 (Fig. 2E). This result suggests that TOBi85 and TOBi89 bind to distinct CEACAM6 epitopes, and that these epitopes may be differentially expressed in SUIT2 cells. Indeed, CEACAM6 is known to be heavily glycosylated, and antigenic variants of this glycoprotein have been previously described [20-23].

To identify the cognate antigens of the 6 non-CEACAM TOBi-bodies, we employed a similar strategy using IP/MS followed by CRISPR/Cas9-mediated ablation of the putative antigens. Accordingly, we found that the cognate antigen of TOBi42, TOBi44, TOBi46, and TOBi117 is ITGA2 (Fig. S5A and S5B, Table S5). The TOBi97 antigen is ITGA3 (Fig. 5C and Table S6), whereas the TOBi65 antigen is MUC1 (Fig. S5D, Table S7).

In summary, we have identified the cognate antigens of all TOBi-bodies (Table S8) and found that the binding specificities of these mAbs converge on antigens from three glycoprotein families, integrins, mucins, and CEACAMs. This suggests that these antigens are highly represented in cell surface repertoire of the PDOs used for immunization and may be broadly expressed across PDAC tissues. As several TOBi-bodies reacted heterogeneously to two members of the CEACAM family, CEACAM5 and CEACAM6, these mAbs may bind to unique cognate epitopes and define variants of these highly glycosylated proteins.

### TOBi85 and TOBi89 Distinguish PDAC from Pancreatitis and Various Normal Tissues

We have demonstrated that the TOBi-bodies bind to PDAC but show little to no reactivity to normal pancreatic tissues. We also identified the cell surface antigens of these TOBi-bodies as ITGA2, ITGA3, CEACAM6, CEACAM5, and MUC1. As these proteins are expressed in various tissues, and antibody binding to normal tissues would preclude the use of these TOBi-bodies for diagnostic imaging, we sought to determine whether these mAbs display binding to tumor-specific epitopes on these cell surface proteins or to normal epitopes expressed in non-pancreatic tissues. To this end, we obtained FFPE tissue microarrays containing 20 types of normal organs and tested the immunoreactivity of the TOBi-bodies and commercially available antibodies raised against peptides from these proteins. Integrin-reactive TOBi-bodies (TOBi42, TOBi44, TOBi46, TOBi97, TOBi117) bind their cognate antigens only on frozen tissues, but we could not obtain frozen tissue microarrays. Therefore, we could only evaluate the reactivity of these mAbs in available pancreas-adjacent frozen tissues such as the duodenum. We found that, with the exception of TOBi65, TOBi85, and TOBi89, the TOBi-bodies generally exhibited binding patterns similar to commercially available antibodies (Table S9), implying that the differential binding of these mAbs in PDAC tumors is mostly due to the elevated expression of their cognate antigens rather than to tumor-specific alterations. Particularly, a commercially available CEACAM6 mAb and most CEACAM6-reactive TOBi-bodies antibodies displayed strong binding in various normal tissues, such as the esophagus, colon and stomach, despite showing no reactivity in the normal pancreas (Fig. 3A, Table S9). In contrast, the CEACAM6 specific TOBi-body, TOBi85 only showed minimal reactivity with salivary gland tissue, and TOBi89 showed no binding to any of the normal tissues tested, despite exhibiting strong binding to PDAC (Fig. 3A, Table S9). These data indicate that TOBi85 and TOBi89 bind to unique CEACAM6 epitopes present in PDAC but absent or expressed at low levels in normal tissues.

**Figure 3.**
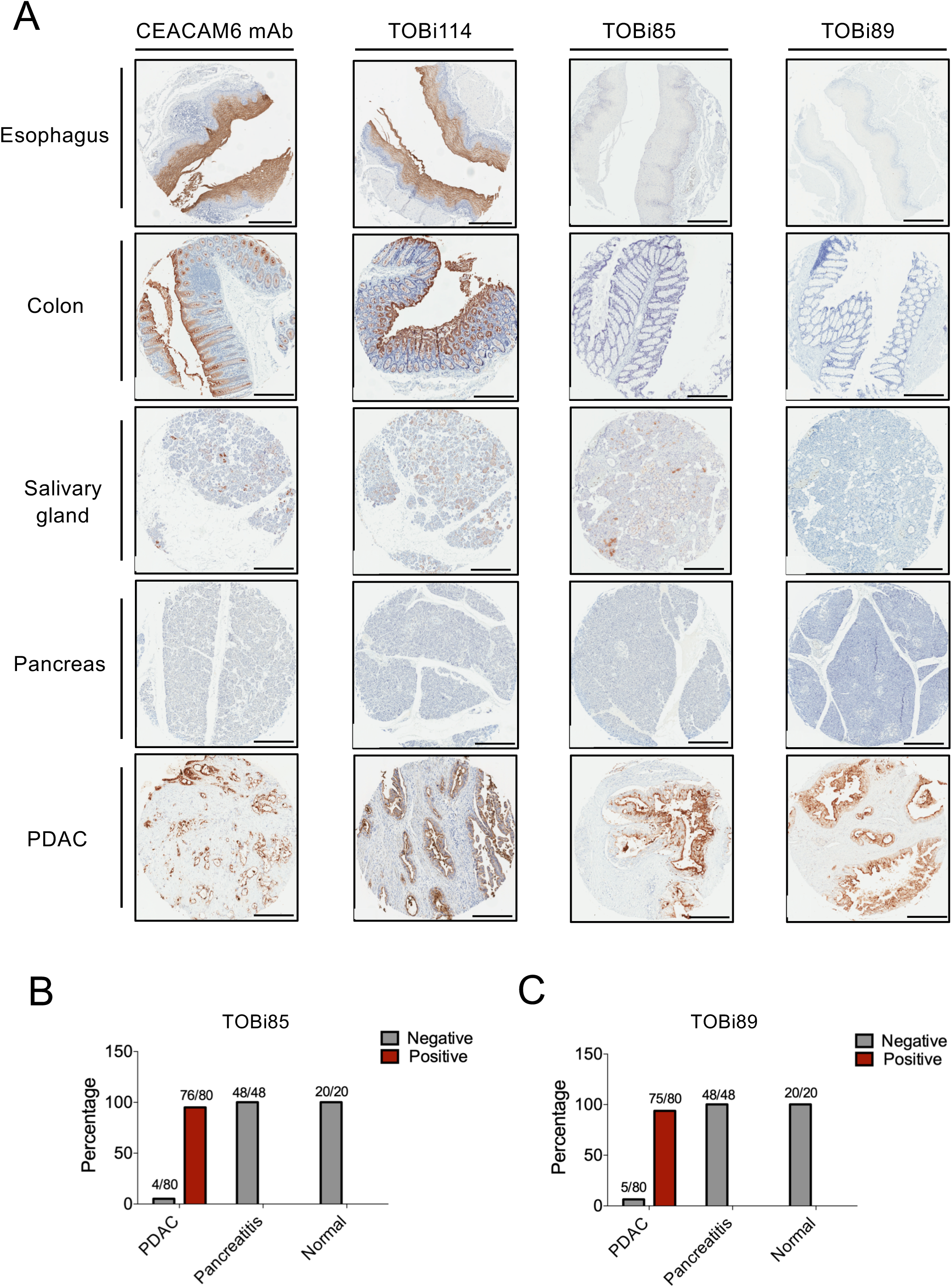
TOBi85 and TOBi89 distinguish PDAC from pancreatitis and various normal tissues (*A*) Immunohistochemical analysis of CEACAM6-specific TOBi-bodies (TOBi85, TOBi89, and TOBi114) and a commercially available CEACAM6 antibody in PDAC and various normal tissues. Antibodies were used for immunohistochemistry in normal tissue microarrays comprising 20 types of organs (3 cores per organ) and in PDAC microarrays comprising 6 PDAC samples. Representative images of antibody immunoreactivity in esophagus, colon, salivary gland, pancreas, and PDAC are shown. Scale bars, 400 μm. (*B* and *C*) TOBi85 and TOBi89 discriminate PDAC from pancreatitis. TOBi85 and TOBi89 were used for immunohistochemistry in a pancreatitis tissue microarray comprising 3 acute and 45 chronic pancreatitis tissue cores, and in a tumor tissue microarray comprising 80 PDAC tissue cores and 20 normal pancreas cores. TOBi85 and TOBi89 displayed reactivity (positive, red bars) in 95% and 93.7% of PDAC tissues respectively, whereas no labeling (negative, gray bars) was observed in all pancreatitis and normal pancreas specimens.

Although TOBi85 and TOBi89 distinguish PDAC from various normal tissues, the clinical utility of these TOBi-bodies for diagnostic immunoPET imaging depends on their ability to detect PDAC in most patients and to distinguish PDAC from benign diseases such as pancreatitis.

Therefore, we assessed the binding of these mAbs in a pancreatitis tissue microarray (48 cores) and a tissue microarray containing 20 normal pancreas controls and 80 tumor tissue cores from patients at various PDAC stages, ranging from stage 1b to 4. Both TOBi85 and TOBi89 showed no binding to normal pancreas or chronic pancreatitis tissues, but reacted with the majority of PDAC tissues (76/80 and 75/80 tumor cores respectively) (Fig. 3B and 3C). Hence, these results validate the broad tumor specificity of these 2 mAbs and are consistent with previous reports of CEACAM6 upregulation in PDAC tissues [24], thereby supporting the development of these antibodies for immunoPET imaging of PDAC.

Overall, we have identified two TOBi-bodies, TOBi85 and TOBi89, with reactivity to epitopes minimally expressed by normal tissues and highly expressed in PDAC. This tumor-specificity is a crucial parameter for the potential diagnostic imaging applications of these TOBi-bodies.

### TOBi85 and TOBi89 Enable PDAC Detection by ImmunoPET Imaging in Mouse Models

To test whether TOBi85 and TOBi89 could detect PDAC *in vivo*, we first conjugated both TOBi-bodies with zirconium-89 (^89^Zr) using the iron chelator, deferoxamine (DFO) (Fig. S6A). We then assessed the purity of ^89^Zr-labeled TOBi85 and TOBi89 by radio instant thin layer chromatography (radio-iTLC). Both radiolabeled mAbs exhibited >99% radiochemical purity, and the specific activity of ^89^Zr-labeled TOBi85 and TOBi89 were 16.7 and 14.3 µCi/µg respectively (Fig. S6B), indicating that there were no significant differences in the quantity of radiolabels attached to these mAbs. Several studies have demonstrated the limited stability of ^89^Zr-based radiolabeled antibodies *in vivo*, resulting in non-specific accumulation to the bone [25, 26]. Hence, we sought to predict the *in vivo* stability of these radiotracers by determining whether they are stable in serum *in vitro*. We incubated ^89^Zr-labeled TOBi85 and TOBi89 with either saline, PBS, and serum at 37°C for 24 hours and then performed radio-iTLC to determine radiochemical purity. We found that both radiolabeled antibodies retained 100% of bound-^89^Zr in serum compared to PBS and saline (Fig. S6C). This result shows that bound-^89^Zr on these radiolabeled antibodies is stable in serum *in vitro* and will likely remain conjugated *in vivo*.

To test the *in vivo* tumor binding of these TOBi-bodies, we first established subcutaneous xenografts of SUIT2 cells in the flanks of athymic nude mice. Two weeks after transplantation when these tumors were approximately 5mm in diameter by caliper measurements, we injected the mice intravenously with either ^89^Zr-TOBi85 or ^89^Zr-TOBi89 (80-120 µCi, 5-9µg) and then performed PET imaging at 1, 2, 3, and 6 days post-injection. We observed a time-dependent increase in tumor uptake in the flanks of these mice (Fig. S6D), indicating that both TOBi-bodies can efficiently target tumors in a physiological setting. As antibodies can exhibit non-specific binding to tissues, we wanted to further characterize the off-target biodistribution of these radiolabeled antibodies *in vivo*. To this end, we harvested the tumors and several organs and measured radiotracer uptake in various organs using radio-iTLC. Both ^89^Zr-TOBi85 and ^89^Zr-TOBi89 demonstrated the highest radiotracer uptake in the tumors and relatively low uptake in other tissues (Fig. S6E). These data corroborate the imaging results and demonstrate that these TOBi-bodies target tumors *in vivo*.

Subcutaneous models lack the desmoplastic features of PDAC. Therefore, the efficient radiotracer uptake in subcutaneous PDAC models may not accurately predict the clinical utility of these TOBi-bodies for imaging PDAC. To assess the tumor targeting of ^89^Zr-TOBi85 and ^89^Zr-TOBi89 in an orthotopic setting, we generated orthotopically grafted organoid (OGO) models, which exhibit a robust desmoplastic reaction and faithfully recapitulates the histopathological features of PDAC [27]. We transplanted hT3 PDAC organoids into the pancreas parenchyma of NOD scid gamma (NSG) mice and then measured tumor size by abdominal ultrasound after 3 weeks (Fig. 4A and S7 A-C). Mice with tumors between 3 to 5 mm in diameter were injected with ^89^Zr-TOBi85, ^89^Zr-TOBi89 or ^89^Zr-IgG2A isotype control. We then obtained PET/CT images at 1, 2, 5, 6, and 9 days post-injection. Consistent with the subcutaneous model, we observed high tumor uptake with both radiolabeled TOBi-bodies, whereas ^89^Zr-IgG2A showed no tumor uptake and was rapidly cleared through the intestines, indicating that the radiotracer uptake in ^89^Zr-TOBi85 and ^89^Zr-TOBi89-injected mice is determined by the specific binding of these TOBi-bodies (Fig. 4B and 4C). In mice injected with ^89^Zr-TOBi85 and ^89^Zr-TOBi89, we observed minimal off-target biodistribution to other tissues, such as the bones, wherein we detected tiny focal signals (Fig. 4B). These could be attributed to free ^89^Zr uncoupled from the radiotracers because ^89^Zr-IgG2A-injected mice also exhibited low bone uptake despite rapid radiotracer clearance (Fig 4C).

**Figure 4.**
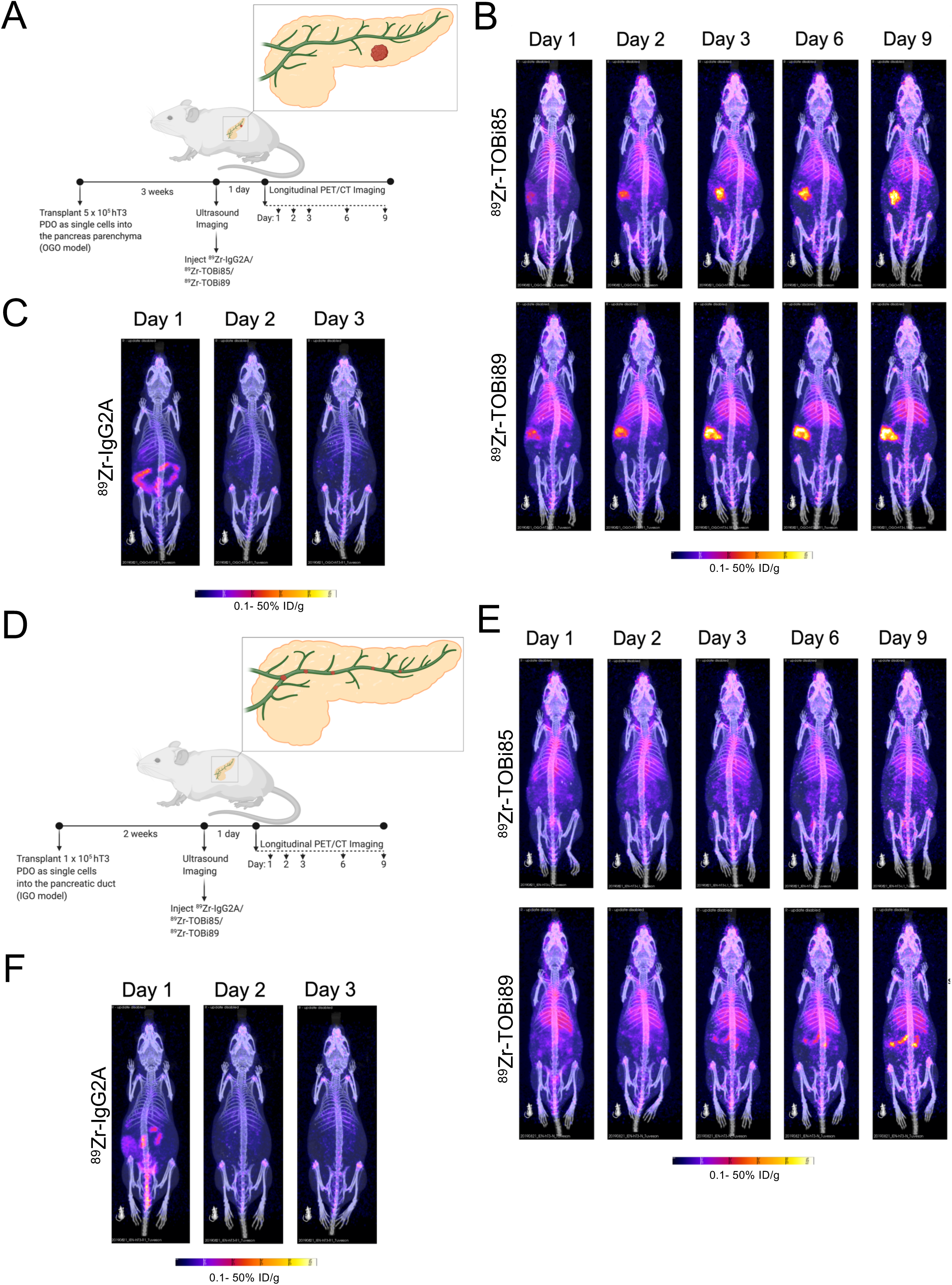
ImmunoPET imaging using TOBi85 and TOBi89 enables the detection of PDAC in mouse models. (*A*) Schematic diagram illustrating the process of immuno-PET imaging in orthotopically-grafted organoid (OGO) transplantation models. Patient-derived organoids (PDOs) are injected into the pancreas parenchyma in the OGO model. (*B*) Serial PET/CT imaging in hT3 OGO models using ^89^Zr-DFO-TOBi85 (top) and ^89^Zr-DFO-TOBi89 (bottom). Maximum image projection is shown. Radiotracer uptake is reported in percentage injected dose per gram of tissue (% ID/g). (*C*) Serial PET/CT imaging in hT3-transplanted OGO models using ^89^Zr-DFO-IgG2A isotype control. Maximum image projection is shown. Radiotracer uptake is reported in percentage injected dose per gram of tissue (% ID/g). (*D*) Schematic diagram illustrating the process of immuno-PET imaging in intraductally-grafted organoid (IGO) transplantation models. PDOs are injected into the pancreas duct in the IGO model and form small (<2 mm) multi-focal tumors along the length of the pancreas. (*E*) Serial PET/CT imaging in hT3 IGO model using ^89^Zr-DFO-IgG2A isotype control. Maximum image projection is shown. Radiotracer uptake is reported in percentage injected dose per gram of tissue (% ID/g). (*F*) Serial PET/CT imaging in hT3 IGO models using ^89^Zr-DFO-TOBi85 (top) and ^89^Zr-DFO-TOBi89 (bottom). Maximum image projection is shown. Radiotracer uptake is reported in percentage injected dose per gram of tissue (% ID/g)

Radiolabeled TOBi85 and TOBi89 detected relatively large tumors, which were also detectable by palpation or high-resolution ultrasound, in the subcutaneous and orthotopic models. However, as detection of small lesions at an earlier stage of disease provides the best chance of survival for PDAC patients [28], we sought to determine the ability of these radiolabeled antibodies to delineate small tumors. To this end, we generated intraductally grafted organoid (IGO) models, which recapitulate PDAC progression from intraductal lesions, contiguous with the pancreatic ductal system, to invasive carcinoma with prominent stromal features [29]. The IGO model is also characterized by small (<2 mm), multi-focal tumors at early stages of intraductal growth. To test these ^89^Zr-labeled TOBi-bodies in the IGO model, we first injected hT3 or hM1a human organoids into the main pancreatic duct of NSG mice and performed abdominal ultrasound after two weeks to detect tumors (Fig. 4D). Unlike in the OGO models, we could not detect any tumors by ultrasound or palpation in the IGO models (Fig. S7D, S7E, S7F, S8A, S8B, and S8C). We then injected the radiotracers and imaged serially with PET/CT as previously done in the OGO models. Although undetectable by abdominal ultrasound, ^89^Zr-TOBi89 delineated multi-focal lesions along the length of the pancreas, consistent with the expected tumor growth pattern in both hT3 and hM1a IGO models (Fig. 4E and S8D), whereas ^89^Zr-TOBi85 exhibited less uptake in hM1a-transplanted IGO model and did not display any evident uptake in the hT3 IGO model (Fig. 4E and S8D). Similar to our observation in the OGO model, ^89^Zr-IgG2A isotype control was rapidly cleared through the liver and intestines (Fig. 4F and S8E). These data, in addition to similar observations in the OGO models, indicate that TOBi89 has a better binding profile to PDAC *in vivo*.

Collectively, our data demonstrate *in vivo* PDAC detection by immunoPET imaging with two tumor-selective TOBi-bodies and show the potential of these mAbs for the diagnostic imaging of PDAC.

## Discussion

The lack of accurate early detection methods underlies the high lethality of PDAC and represents a principal barrier to improving patient outcomes. Here, we sought to address this unmet need by developing novel tumor-specific antibodies and evaluating their potential for detecting incipient PDAC using the highly sensitive immuno-PET modality. Accordingly, we established a pipeline to generate rat-derived mAbs against cell-surface proteins of patient-derived organoids and discovered 16 antibodies with reactivity to PDAC tumors but not matched adjacent normal pancreas. We determined the cognate antigens of these mAbs and identified two CEACAM6-reactive antibodies that exhibit strong binding to PDAC but show minimal binding to various normal tissues and chronic pancreatitis specimens. PET imaging using these mAbs enabled the detection of human organoid-derived tumors that were otherwise undetectable by high resolution ultrasound imaging techniques in PDAC mouse models. These results indicate the potential application of these mAbs for facilitating the early detection of resectable PDAC and, thereby, prolonging patient survival.

Previous studies have used CEACAM6 antibodies for immunoPET imaging of large subcutaneous tumors [30, 31]. Here, we identify two mAbs, TOBi85 and TOBi89, that bind to tumor-specific variants of CEACAM6. We also demonstrate the potential of these TOBi-bodies for detecting both large and small tumors in orthotopic mouse models that faithfully recapitulate PDAC progression [27, 29]. Indeed, both radiolabeled antibodies detected 3-5mm tumors in OGO models, whereas ^89^Zr-TOBi89 consistently delineated <2mm multi-focal lesions in the IGO model. However, we did not observe strong radiotracer signal until day 5 for small tumors in this model. This slow accumulation, in addition to long clearance kinetics, are well-established limitations of immunoPET imaging using ^89^Zr-labeled full-size antibodies [32]. This contrasts with FDG-PET imaging, which typically peaks within few hours [33], resulting in low patient exposure to radiation and allowing for feasible longitudinal surveillance studies. This limitation could be addressed in three ways. First, the generation of smaller derivatives of these TOBi-bodies, such as antigen-binding fragments (Fab) or single-chain variable fragments (scFv), could facilitate increased tissue penetration, faster pharmacokinetics, and enable the use of shorter-lived radionuclides for imaging. Second, pre-targeted immunoPET approaches, wherein an unlabeled antibody is injected and radiolabeling of the antibody occurs *in vivo*, will drastically reduce patient exposure to radiation by uncoupling the pharmacokinetics of the antibody from the decay properties of the PET radioisotope [34]. Third, the use of highly sensitive PET systems, such as the recently developed total-body PET system with 40 times higher sensitivity [35], will reduce patient radiation exposure since these systems require lower radiotracer concentration.

We demonstrate that TOBi85 and TOBi89 discriminate PDAC from chronic pancreatitis, a benign pancreatic disease and common confounder in the diagnosis of PDAC [36, 37]. Therefore, these mAbs would enable accurate identification of incipient PDAC. As these antibodies bind to CEACAM6, and high gene expression of this glycoprotein has been associated with the classical subtype of PDAC [38], the clinical utility of these TOBi-bodies could be limited to patients with classical PDAC. However, we show here that these antibodies bind to the majority of PDAC tissues, consistent with prior reports of CEACAM6 antibody reactivity [24]. In addition, the IGO model enables the modeling of classical and basal subtypes in vivo, and we successfully transplanted and detected radiotracer uptake in engrafted human organoids, hT3 and hM1a, which display basal-like subtype *in vivo* [29]. Therefore, immunoPET imaging using these CEACAM6-reactive TOBi-bodies can be broadly applicable in PDAC.

Here, we employed mouse models to demonstrate the potential of TOBi85 and TOBi89 for immunoPET imaging. We previously demonstrated that both TOBi-bodies bind to CEACAM6 epitopes that are not expressed in various human tissues. We also established that radiolabeled TOBi85 and TOBi89 exhibited low off-target biodistribution in mouse tissues. Therefore, we would expect a minimal accumulation of these radiolabeled TOBi-bodies in non-malignant tissues in human patients. Since these studies were performed in immuno-compromised mice, thereby excluding the contribution of the acquired immune system to the pharmacokinetic parameters, and mice do not express CEACAM6, accurate assessment of the off-target and on-target biodistributions of these radiotracers will require imaging studies in human subjects.

The clinical translation of these rat-derived antibodies will require humanization of the antibody Fc and framework regions because of the well-documented immunogenicity of rodent antibodies in humans [39]. Additionally, the immunoPET modality is expensive and not widely available. Thus, accurate non-invasive serological tests are still required to enrich for at-risk groups that will maximally benefit from highly sensitive and specific immunoPET/CT diagnostic imaging. In future studies, we will evaluate the utility of these TOBi-bodies for the serological detection of incipient PDAC. Additionally, we will explore the therapeutic potentials of these mAbs as antibody-drug conjugates, and for antibody-targeted radiotherapy and immunotherapy.

In conclusion, we integrated the tumor-specificity of two CEACAM6 antibodies, TOBi85 and TOBi89, with the high sensitivity of PET imaging to detect tumors in several transplantation mouse models of PDAC. The data presented herein support the clinical development of these antibodies as non-invasive PET radiotracers for facilitating the staging and early detection of PDAC.

## Methods

### Mouse Models

NOD scid gamma (NSG, stock number 005557) mice were purchased from The Jackson Laboratory; *nu/nu* mice (stock number 24102242) were purchased from The Charles River Laboratory. All animal procedures and studies were conducted in accordance with the Institutional Animal Care and Use Committee at Cold Spring Harbor Laboratory (CSHL) and Memorial Sloan Kettering Cancer Center (MSKCC).

### *In Vivo* Transplantations

Injections for the generation of subcutaneous tumors were performed as previously described [40]. 1 × 10^6^ SUIT2 cells were injected into the flanks of nude mice. Injections for the generation of orthotopically-grafted organoid (OGO) tumors were conducted as previously described [27]. 5 × 10^5^ single cells prepared from organoid cultures were resuspended as a 45 μL suspension of 50% Matrigel in PBS and injected into the pancreas parenchyma. For intraductal grafting of organoid (IGO) transplants, 100,000 cells, suspended in 25 μl of PBS, were infused into the pancreatic duct using the modified retrograde infusion technique described previously [29, 41].

### Generation of Rat Hybridomas

Patient derived-organoids (PDOs) were harvested using cell recovery solution (Corning cat. no. 354253) and then pooled. The organoids were then washed with 10 mL ice-cold PBS. PBS was completely removed and 3 ml of Digitonin solution (42 µg/ml digitonin, 2 mM DTT, 2 mM MgCl_2_, 150 mM NaCl, 0.2 mM EDTA, 20 mM HEPES pH7.4) was added. The solution was gently mixed for 10 min at 4°C, and then pelleted by centrifugation at 800 g for 5 min. The supernatant containing cytoplasmic fractions was discarded, and the pellet was homogenized in 200 µL 1x ice-cold PBS by passing through a 23-gauge needle. Three 6-week-old Sprague Dawley rats (Taconics) were immunized with PDO membrane fractions (100 mg per animal per boost, for five weekly boosts). Immune response was monitored by Cell-based ELISA using SUIT2 cells to measure the anti-PDAC serum IgG titer. After a 60-day immunization course, the rats were harvested and 10^8^ splenocytes were collected for making hybridomas by fusing with the rat myeloma cell line YB2/0, following the standard method [42]. All procedures were approved by the Cold Spring Harbor Laboratory Institutional Animal Care and Use Committee.

### Cell Lines and Cell Culture

Human pancreatic organoid lines were cultured as previously described [27]. All human organoid experiments were reviewed and approved by the Institutional Review Board of Cold Spring Harbor Laboratory, and conducted in accordance with recognized ethical guidelines (Declaration of Helsinki). All cells were cultured at 37°C with 5% CO_2_. Cell line authentication was not performed. *Mycoplasma* testing with the MycoAlert Mycoplasma Detection Kit (LT07-318; Lonza) is performed monthly at our institution, and each cell line has been tested at least once after thawing or isolation, and retested prior to the transplantation experiments.

### Cell-based ELISA

SUIT2 cells were seeded at a density of 5 x 10^4^ cells per well in a 100 µL DMEM/10% FBS media volume on 96-well clear flat-bottom tissue culture plates (Corning, cat. no. 3595). After overnight incubation at 37°C, 5% CO_2_ to allow for cell adherence, cells were washed twice with 200 µL cold PBS and then fixed with 100 µL 4% PFA for 10 mins at room temperature. The fixed cells were washed three times with 200 µL PBS and then blocked for 1 hr with 3% BSA in PBS. 100 µL of diluted antibodies or hybridoma supernatant were then added. After a 2 hr incubation at room temperature, cells were washed four times with 200 µL PBS and then incubated with 100 µL diluted secondary antibody for 1 hr. Cells were washed four times with 200 µL PBS and incubated with 100 µL TMB substrate (Thermo Sci, cat. No. 34028) for 20 min. 100 µL 1 M phosphoric acid was added to stop the peroxidase reaction and the absorbance measurements were taken at Abs=450 nm using SpectaMax I3 (Molecular Devices).

### Cell-surface Flow Cytometry

Single cell suspensions of cell lines or organoids were prepared. Next, the cells were washed with ice-cold PBS and blocked for 30 mins with 3% BSA in PBS (staining buffer). Primary antibodies were then added to the cells followed by a 30 min incubation. Afterwards, the cells were washed with staining buffer and the appropriate fluorescent secondary antibodies were mixed with DAPI and added to cells. After incubating for 10 min, the cells were washed with staining buffer and analyzed using a LSR Fortessa cytometer (BD Biosciences). Data files were analyzed using FlowJo v10 software (FlowJo, LLC).

### Solid-phase ELISA

CEACAM6 (Sino Biological, cat. no. 10823-H08H) or CA19-9 Antigen (myBiosource, cat. no. MBS318287) recombinant protein diluted in PBS or SUIT2 conditioned media was immobilized overnight on high binding ELISA 96-well microplates (Corning, cat. no. 9018). The wells were then washed three times with 200 µL PBS with 0.05% Tween-20 (PBS-T) and blocked for 2 hr with 200 µL 3% BSA in PBS-T. Diluted antibodies were then added and incubated for 1 hr. Next, the plates were washed three times with PBS-T and then incubated with 100 µL diluted secondary antibody for 1 hr. Afterwards, the wells were washed four times with 200 µL PBS and incubated with 100 µL TMB substrate (Thermo Sci, cat. No. 34028) for 20 min. 100 µL 1 M phosphoric acid was added to stop the peroxidase reaction and the absorbance measurements were taken at Abs=450 nm using SpectaMax I3 (Molecular Devices).

### CRISPR/Cas9 Knockout

Lenti-Cas9-puromycin plasmids were used. Cell lines were infected and selected using 2.5 μg/mL puromycin (A1113803; Thermo Scientific). Single guide RNAs (sgRNA) were designed using CRISPR Design (http://crispr.mit.edu) and cloned into the LRGN (Lenti-sgRNA-EFS-GFP-neo) plasmid. Organoids were infected and plated as single cells in the presence of 2 mg/mL neomycin (10131035; Invitrogen). Knockout was confirmed by Sanger sequencing (data not shown), western blot (data not shown), and flow cytometric analysis.

### Human Specimens

Human PDAC resection specimens were obtained from the Sidney Kimmel Comprehensive Cancer Center at Johns Hopkins University and from Memorial Sloan Kettering Cancer Center. All tissue donations and experiments were reviewed and approved by the Institutional Review Board of Cold Spring Harbor Laboratory and the clinical institutions involved. Written informed consent was obtained prior to acquisition of tissue from all patients. The studies were conducted in accordance with ethical guidelines (Declaration of Helsinki). Tissue microarrays were obtained from US Biomax (BBS14011, PA1001a, PA241d, FDA999q).

### Immunohistochemical Labeling

Standard procedures were used for immunohistochemistry (IHC) with the TOBi-bodies, and CEACAM6 (Millipore Sigma, cat. no. MABT323), MUC1 (GeneTex, cat. no. GTX10114), ITGA2 (Sigma, cat. no. HPA063556), and ITGA3 (Atlas, cat. no. HPA008572) antibodies.

### Ultrasound Imaging

Ultrasound imaging of OGO and IGO tumors was performed using the Vevo 3100 Visual Sonics Preclinical Imaging System (Visualsonics, Canada) with an MX550D ultra-high frequency linear array transducer, which enables the detection of tumors between 1-30 mm in diameter. To perform the scan, mice were anesthetized with 3% of isoflurane and oxygen. Each mouse was kept warm using a heated platform while scanning. Warm Aqausonic gels were also placed at the right lateral side of the mice. Vevo Lab Software analysis was used to analyze the images and perform annotations and measurements.

### Radiolabeling Antibodies with ^Zr^89

Radiolabeling of Rat IgG2A (BioXcell, cat. no. BE0089), TOBi85 and TOBi89 with ^89^Zr-DFO was performed as previously described [43-45]. The ^89^Zr-oxalate in oxalic acid (1 M) was neutralized to pH 7.0–7.2, using Na_2_CO_3_ (1 M) followed by addition of the appropriate construct in PBS (pH 7.4). The mixture was incubated at room temperature for 1–2 h and monitored using radio-iTLC, eluting with an aqueous solution of EDTA (50 mM, pH 5.5). After adequate radiolabeling, the reaction was quenched by addition of the same EDTA solution and the labeled construct was purified using gel-filtration chromatography (Sephadex G-25, PD10 desalting column; GE Healthcare) into 0.9% saline. Radiochemical purity was assessed by iTLC in an aqueous solution of EDTA (50 mM, pH5.5). The ^89^Zr is produced at MSKCC via the ^89^Y(*p,n*)^89^Zr transmutation reaction on a TR19/9 variable-beam energy cyclotron (Ebco Industries) [44]. All activity measurements were performed using a Capintec CRC-15R dose calibrator. Instant TLC (ITLC) for radio-iTLC experiments was performed on strips of glass-fiber, silica-impregnated paper (PallCorp.), read on a Bioscan AR-2000 radioTLC plate reader, and analyzed using Winscan Radio-TLC software (Bioscan).

### PET and PET/CT Imaging

Mice were administered 89Zr-labeled antibodies (80-120 μCi, 5-9μg) via tail vein injection and images were acquired at day 1, 2, 5, 6, and 9. For PET imaging in subcutaneous models, imaging was performed at day 1, 2, 3, and 6. Static scans were recorded on the PET/CT scanner at the various time points with a minimum of 30 million coincident events (10–30 min total scan time). Data were sorted into two-dimensional histograms by Fourier rebinning, and the images were reconstructed using a 2D ordered subset expectation-maximization (2DOSEM) algorithm (16 subsets, four iterations) into a 128 × 128 × 159 (0.78 × 0.78 × 0.80 mm) matrix. The image data were normalized to correct for nonuniformity of response of the PET, dead-time count losses, positron branching ratio, and physical decay to the time of injection but no attenuation, scatter, or partial-volume averaging correction was applied. Activity concentrations [percentage of dose per gram of tissue (% ID/g) and MIPs were determined by conversion of the counting rates from the reconstructed images. Whole-body CT scans were acquired with a voltage of 80 kV and 500 μA. A total of 120 rotational steps for a total of 220° were acquired with a total scan time of 120 s and 145 ms per frame exposure. Combined PET/CT images were processed and optimized to show localization of the PET signal in the tumor.

### Statistical Analysis and Schematic diagrams

GraphPad Prism was used for graphical representation of data. Statistical analysis was performed using Student *t* test. Schematic diagrams were created using Biorender.com.

### Tryptic Digestion and iTRAQ Labeling

The beads were reconstituted with 20 µL of 50 mM triethylammonium bicarbonate buffer (TEAB). RapiGest was added to a final concentration of 0.1% and *tris*(2-carboxyethyl)phosphine (TCEP) was added to final concentration of 5 mM. Samples were then heated to 55°C for 20 min, allowed to cool to room temperature and methyl methanethiosulfonate (MMTS) added to a final concentration of 10 mM. Samples were incubated at room temperature for 20 min to complete blocking of free sulfhydryl groups. 2 µg of sequencing grade trypsin (Promega) was then added to the samples and they were digested overnight at 37°C. After digestion the supernatant was removed from the beads and was dried in vacuo. Peptides were reconstituted in 50 µL of 0.5 M TEAB/70% ethanol and labeled with 8-plex iTRAQ reagent for 1 hour at room temperature essentially according to Ross et al [46]. Labeled samples were then acidified to pH 4 using formic acid, combined and concentrated in vacuum until ∼10 µL remained.

### 2-Dimensional Fractionation

Peptides were fractionated using a Pierce High pH Reversed-Phase Peptide Fractionation Kit (Thermo Scientific) according to the manufacturer’s instructions with slight modifications. Briefly, peptides were reconstituted in 150 µL of 0.1% TFA, loaded onto the spin column and centrifuged at 3000 g for 2 min. Column was washed with water and then peptides were eluted with the following percentages of acetonitrile (ACN) in 0.1% triethylalmine (TEA): 5%, 7.5%, 10%, 12.5%, 15%, 20%, 30% and 50%. Each of the 8 fractions was then separately injected into the mass spectrometer using capillary reverse phase LC at low pH.

### Mass Spectrometry

An Orbitrap Fusion Lumos mass spectrometer (Thermo Scientific), equipped with a nano-ion spray source was coupled to an EASY-nLC 1200 system (Thermo Scientific). The LC system was configured with a self-pack PicoFrit™ 75-μm analytical column with an 8-μm emitter (New Objective, Woburn, MA) packed to 25 cm with ReproSil-Pur C18-AQ, 1.9 µM material (Dr. Maish GmbH). Mobile phase A consisted of 2% acetonitrile; 0.1% formic acid and mobile phase B consisted of 90% acetonitrile; 0.1% formic Acid. Peptides were then separated using the following steps: at a flow rate of 200 nL/min: 2% B to 6% B over 1 min, 6% B to 30% B over 84 min, 30% B to 60% B over 9 min, 60% B to 90% B over 1 min, held at 90% B for 5 min, 90% B to 50% B over 1 min and then flow rate was increased to 500 nL/min as 50% B was held for 9 min. Eluted peptides were directly electrosprayed into the Fusion Lumos mass spectrometer with the application of a distal 2.3 kV spray voltage and a capillary temperature of 300°C. Full-scan mass spectrum (Res=60,000; 400-1600 *m/z*) were followed by MS/MS using the “Top Speed” method for selection. High-energy collisional dissociation (HCD) was used with the normalized collision energy set to 35 for fragmentation, the isolation width set to 1.2 and a duration of 10 sec was set for the dynamic exclusion with an exclusion mass width of 10 ppm. We used monoisotopic precursor selection for charge states 2+ and greater, and all data were acquired in profile mode.

### Database Searching

Peaklist files were generated by Mascot Distiller (Matrix Science). Protein identification and quantification was carried using Mascot 2.4 [47] against the UniProt human sequence database (92,919 sequences; 36,868,442 residues). Methylthiolation of cysteine and N-terminal and lysine iTRAQ modifications were set as fixed modifications, methionine oxidation and deamidation (NQ) as variable. Trypsin was used as cleavage enzyme with one missed cleavage allowed. Mass tolerance was set at 30 ppm for intact peptide mass and 0.3 Da for fragment ions. Search results were rescored to give a final 1% FDR using a randomized version of the same Uniprot Human database, and at least two scoring peptides were required. Protein-level iTRAQ ratios were calculated as intensity weighted, using only peptides with expectation values < 0.05. As this was a protein IP experiment, no global ratio normalization was applied. Protein enrichment was then calculating by dividing the true sample protein ratios by the corresponding control sample ratios.

## Acknowledgements

We acknowledge the Cold Spring Harbor Laboratory Antibody and Phage Display, Animal and Genetic Engineering, Animal and Tissue Imaging, Mass Spectrometry, Flow Cytometry and Microscopy Shared Resources, which are supported by the NIH Cancer Center Support Grant P30CA045508. The authors are supported by the NIH Cancer Center Support Grant 5P30CA045508 and the Lustgarten Foundation, where D.A. Tuveson is a distinguished scholar and Director of the Lustgarten Foundation–designated Laboratory of Pancreatic Cancer Research. D.A. Tuveson is also supported by the Thompson Foundation, the Cold Spring Harbor Laboratory and Northwell Health Affiliation, the Northwell Health Tissue Donation Program the Cold Spring Harbor Laboratory Association and the National Institutes of Health (NIH 5P30CA45508, U01CA210240, R01CA229699, U01CA224013, 1R01CA188134, and 1R01CA190092). This work was also supported by a gift from the Simons Foundation (552716 to D.A. Tuveson), and by a gift from the Wojcicki-Troper Foundation (to R.H. Hruban). J.S. Lewis is supported by the NIH grant, NCI R35 CA232130. G. Biffi was a fellow of the Human Frontiers Science Program (LT000195/2015-L) and EMBO (ALTF 1203-2014).

## Authors’ contribution

Conceptualization: T.E. Oni, J.T. Yeh, S.K. Lyons, D.A. Tuveson

Investigation: T.E. Oni, C. Bautista, J. Merrill, J. Goos, K.D. Rivera, K. Miyabayashi, G. Biffi, L. Garcia, M. Samaritano

Methodology: T.E. Oni, J.S. Lewis, D.J. Pappin, S.K. Lyons, J.T. Yeh

Formal analysis and interpretation of data: T.E. Oni, J. Merrill, J. Goos, L. Garcia, K.D. Rivera, D.J. Pappin

Visualization: T.E. Oni, J. Merrill, J. Goos

Writing – original draft: T.E. Oni

Writing – review & editing: T.E. Oni, D.A. Tuveson, G. Biffi, R.H. Hruban, J.S. Lewis, SK. Lyons

Project administration: T.E. Oni

Resources: D. Plenker, H. Patel, E. Elyada, K. Yu, M. G. Goggins, R. H. Hruban

Supervision: D.A. Tuveson, J.S. Lewis, S.K. Lyons, J.T. Yeh

Funding acquisition: D.A. Tuveson, J.S. Lewis, S.K. Lyons, J.T. Yeh

## Figure Legends

**Supplementary Figure S1**. Antibody screening process. (*A*) Cell-based ELISA schematic for screening antibodies secreted from rat hybridoma clones. (*B*) Cell-based ELISA determination of antibody responses to the human pancreatic cancer cell line SUIT2. The three rats immunized with membrane fractions of patient-derived organoids (red) displayed high serum IgG antibody titers compared to 2 non-immunized animals (blue). Data shown as mean of duplicate wells. (*C*) Cell-based ELISA screen of secreted antibodies from hybridoma clones on SUIT2 PDAC cells. Relative absorbance was determined by the ratio of Abs_450nm_ (absorbance at 450 nm) of antibody clones to rat IgG control. A relative absorbance cutoff of 1.5 separates non-binding or negative clones (black circles) from binding or positive clones (red circles). (*D*) Solid-phase ELISA to determine CA19-9-binding antibody clones. Relative absorbance was determined by the ratio of Abs_450nm_ (absorbance at 450 nm) of antibody clones to rat IgG control. 12 out of 77 candidate antibodies displayed binding to CA19-9 (red circles) comparable to NS19-9, an anti-CA19-9 control (blue circle). 65 non-CA19-9 clones (black circles) may contain antibodies with binding specificities to novel tumor epitopes. Data shown as mean of triplicate wells.

**Supplementary Figure S2**. Candidate antibodies (red) display cell-surface binding compared to rat IgG control (blue) in SUIT2 PDAC cells by cell-surface flow cytometric analysis. Results are representative of three independent experiments. FSC-A, forward scatter-area.

**Supplementary Figure S3**. Candidate antibodies (red) display cell-surface binding compared to rat IgG control (blue) in SUIT2 PDAC cells by cell-surface flow cytometric analysis. Results are representative of three independent experiments. FSC-A, forward scatter-area.

**Supplementary Figure S4**. Identification of tumor-selective TOBi-bodies. Immunohistochemical labeling of human PDAC (left) and adjacent normal tissue sections (right) with a panel of TOBi-bodies. Inserts: magnifications. Scale bars, 100 μm.

**Supplementary Figure S5**. Identifying the cognate antigens of the TOBi-bodies. CRISPR knockout of putative antigens, ITGA2, ITGA3, and MUC1 were independently generated in SUIT2 cells. The various TOBi-bodies were tested by flow cytometric analysis for cell surface binding in SUIT2 cells that are either wildtype (WT, red) or knockout (KO, blue) for the respective antigens. (*A*) Validation of ITGA2-specific ablation in SUIT2 cells by CRISPR. Knockout of ITGA2 abolishes the binding of an ITGA2 antibody, but does not reduce the binding of an ITGB1 antibody. (*B*) Knockout of ITGA2 in SUIT2 cells abrogates the binding of TOBi42, TOBi44, TOBi46, and TOBi117. (*C*) Validation of ITGA3 knockout using a commercially available antibody (left panel). Knockout of ITGA3 abrogates the binding of TOBi97 (right panel). (*D*) Validation of MUC1 knockout using a commercially available antibody (left panel). Knockout of MUC1 abrogates the binding of TOBi65 (right panel). Results are representative of three independent experiments. FSC-A, forward scatter-area.

**Supplementary Figure S6**. ImmunoPET imaging using TOBi85 and TOBi89 in subcutaneous mouse models. (*A*) Schematic diagram illustrating the conjugation of the TOBi-bodies with zirconium-89 using the iron chelator, deferoxamine. (*B*) Radiolabeled TOBi-bodies exhibit high radiochemical purity. Radio iTLC analysis of ^89^Zr-TOBi85 and ^89^Zr-TOBi89 shows a major single peak corresponding to radiolabeled antibodies. (*C*) Serum stability of radiolabeled TOBi-bodies. Radio-iTLC analysis of ^89^Zr-TOBi85 (left) and ^89^Zr-TOBi89 (right) shows that these antibodies mostly remain conjugated to the radionuclide and retain radioactivity after 24 hours incubation in serum, saline, or PBS. (*D*) Serial PET imaging in SUIT2-transplanted subcutaneous models using ^89^Zr-DFO-TOBi85 (top) and ^89^Zr-DFO-TOBi89 (bottom). Maximum image projection is shown. Radiotracer uptake is reported in percentage injected dose per gram of tissue (% ID/g). (*E*) Radiotracer uptake of radiolabeled TOBi-bodies in mouse tissues. Biodistribution of ^89^Zr-TOBi85 (white bars) and ^89^Zr-TOBi89 (black bars) in various tissues from SUIT2-transplanted subcutaneous models. Radiotracer uptake is reported in percentage injected dose per gram of tissue (% ID/g). Most radiotracer uptake was in the tumors.

**Supplementary Figure S7**. Ultrasound imaging in orthotopic mouse models. (*A-C*) hT3-transplanted orthotopically-grafted organoids (OGO) mice were evaluated by abdominal ultrasound. Tumors and adjacent tissues are annotated. Mouse (*A*) was assigned to ^89^Zr-TOBi85, (*B*) to ^89^Zr-TOBi89, and (*C*) to ^89^Zr-IgG2a isotype control for immunoPET imaging. (*D-F*) hT3-transplanted intraductally-grafted organoid (IGO) mice were evaluated by abdominal ultrasound. No tumors were evident by this method. The pancreas and adjacent tissues are annotated. Mouse (*D*) was assigned to ^89^Zr-TOBi85, (*E*) to ^89^Zr-TOBi89, and (*F*) to ^89^Zr-IgG2a isotype control for immunoPET imaging.

**Supplementary Figure S8**. ImmunoPET imaging using TOBi85 and TOBi89 in hM1a-transplanted intraductally-grafted organoid (IGO) models (*A-C*) hM1a-transplanted intraductally-grafted organoids (IGO) mice were first evaluated by abdominal ultrasound. No tumors were evident by this method. The pancreas and adjacent tissues are annotated. Mouse (*A*) was assigned to ^89^Zr-TOBi85, (*B*) was assigned to ^89^Zr-TOBi89, and (*C*) to ^89^Zr-IgG2a isotype control for immunoPET imaging. (*D*) Serial PET imaging in hM1a-transplanted IGO models using ^89^Zr-DFO-TOBi85 (top) and ^89^Zr-DFO-TOBi89 (bottom). Maximum image projection is shown. Radiotracer uptake is reported in percentage injected dose per gram of tissue (% ID/g). (*E*) Serial PET/CT imaging in hM1a-transplanted IGO model using ^89^Zr-DFO-IgG2A isotype control. Maximum image projection is shown. Radiotracer uptake is reported in percentage injected dose per gram of tissue (% ID/g).

**Supplementary Table 1**. Panel of patient-derived organoids used for immunization. hT denotes organoids derived from resected samples, and hF denotes organoids obtained from fine needle aspirates. Additional patient information include sex, age, race, tumor stage and treatment status at the time samples were obtained. The genetic status of KRAS and TP53, the predominantly altered oncogene and tumor suppressor respectively, is also indicated.

**Supplementary Table 2**. Immunoprecipitation and mass spectrometry (IP/MS) identify CEACAM6 and CEACAM5 as the likely antigen of TOBi109, TOBi111, and TOBi125. Isobaric tags for relative and absolute quantitation (iTRAQ) mass spectrometry was used to determine the abundance of proteins immunoprecipitated by TOBi109, TOBi111, TOBi125 or rat IgG in SUIT2 cells. Enrichment was determined by calculating ratios of protein abundance in TOBi109, TOBi111, or TOBi125 relative to IgG-precipitated samples and ranked from the highest to lowest enriched proteins. Non-enriched proteins (with enrichment below 1) were filtered out, and an enrichment cut-off of 1.5 was applied (bolded). Subcellular location of enriched proteins was noted based on UNIPROT annotations, and membrane proteins were prioritized. CEACAM6 and CEACAM5 (highlighted) are the top membrane proteins enriched by all three TOBi-bodies.

**Supplementary Table 3**. Immunoprecipitation and mass spectrometry (IP/MS) identify CEACAM6 as the likely antigen of TOBi85. Isobaric tags for relative and absolute quantitation (iTRAQ) mass spectrometry was used to determine the abundance of proteins immunoprecipitated by TOBi85 or rat IgG in BXPC3 and MIAPACA2 cells. Enrichment was determined by calculating ratios of protein abundance in TOBi85 relative to IgG-precipitated samples and ranked from the highest to lowest enriched proteins. Non-enriched proteins (with enrichment below 1) were filtered out, and an enrichment cut-off of 1.5 was applied (bolded). Subcellular location of enriched proteins was noted based on UNIPROT annotations, and membrane proteins were prioritized. CEACAM6 (highlighted) is the only membrane protein enriched in both BXPC3 and MIAPACA2 cells.

**Supplementary Table 4**. Immunoprecipitation and mass spectrometry (IP/MS) identify CEACAM6 as the likely antigen of TOBi89. Isobaric tags for relative and absolute quantitation (iTRAQ) mass spectrometry was used to determine the abundance of proteins immunoprecipitated by TOBi89 or rat IgG in hM19D patient-derived organoids (PDOs). Enrichment was determined by calculating ratios of protein abundance in TOBi89 relative to IgG-precipitated samples and ranked from the highest to lowest enriched proteins. Non-enriched proteins (with enrichment below 1) were filtered out, and an enrichment cut-off of 1.5 was applied (bolded). Subcellular location of enriched proteins was noted based on UNIPROT annotations, and membrane proteins were prioritized. CEACAM6 (highlighted) is the only membrane protein enriched in hM19D PDOs.

**Supplementary Table 5**. Immunoprecipitation and mass spectrometry (IP/MS) identify ITGA2 as the likely antigen of TOBi44 and TOBi46. Isobaric tags for relative and absolute quantitation (iTRAQ) mass spectrometry was used to determine the abundance of proteins immunoprecipitated by TOBi44 and TOBi46 or rat IgG in SUIT2 cells. Enrichment was determined by calculating ratios of protein abundance in TOBi44 and TOBi46 relative to IgG-precipitated samples and ranked from the highest to lowest enriched proteins. Non-enriched proteins (with enrichment below 1) were filtered out, and an enrichment cut-off of 1.5 was applied (bolded). Subcellular location of enriched proteins was noted based on UNIPROT annotations, and membrane proteins were prioritized. ITGA2 (highlighted) is the top membrane protein enriched by both TOBi-bodies.

**Supplementary Table 6**. Immunoprecipitation and mass spectrometry (IP/MS) identify ITGA3 as the likely antigen of TOBi97. Isobaric tags for relative and absolute quantitation (iTRAQ) mass spectrometry was used to determine the abundance of proteins immunoprecipitated by TOBi97 or rat IgG in SUIT2 cells. Enrichment was determined by calculating ratios of protein abundance in TOBi97 relative to IgG-precipitated samples and ranked from the highest to lowest enriched proteins. Non-enriched proteins (with enrichment below 1) were filtered out, and an enrichment cut-off of 1.5 was applied (bolded). Subcellular location of enriched proteins was noted based on UNIPROT annotations, and membrane proteins were prioritized. ITGA3 (highlighted) and its known interactor, CD151 are the top membrane protein enriched by TOBi97

**Supplementary Table 7**. Immunoprecipitation and mass spectrometry (IP/MS) identify MUC1 as the likely antigen of TOBi65. Isobaric tags for relative and absolute quantitation (iTRAQ) mass spectrometry was used to determine the abundance of proteins immunoprecipitated by TOBi65 or rat IgG in hF24 and hF27 patient-derived organoids (PDOs). Enrichment was determined by calculating ratios of protein abundance in TOBi65 relative to IgG-precipitated samples and ranked from the highest to lowest enriched proteins. Non-enriched proteins (with enrichment below 1) were filtered out, and an enrichment cut-off of 1.5 was applied (bolded). Subcellular location of enriched proteins was noted based on UNIPROT annotations, and membrane proteins were prioritized. MUC1 (highlighted) is the top membrane protein enriched in both hF24 and hF27 PDOs.

**Supplementary Table 8**. Validated list of the cognate antigens of the TOBi-bodies.

**Supplementary Table 9**. TOBi85 and TOBi89 display minimal binding to normal human tissues. Immunostaining scores of a panel of TOBi-bodies and commercially-available antibodies raised against the cognate antigens of the TOBi-bodies in normal tissue microarrays and pancreas-adjacent frozen tissues. Scores were assigned 1-5 (low to high) based on the intensity of the staining and highlighted accordingly. Only TOBi65, 85, and 89 deviated significantly from the staining pattern of commercially-available antibodies raised against their cognate antigens. “N/A” denotes that we could not properly assess TOBi44, TOBi42, TOBi44, TOBi46, and TOBi117 as they do not bind to FFPE tissues. However, we noted that they display high binding to pancreas-adjacent small intestines on frozen sections. Scores are representative of three tissue cores or slides.

## References

1. Siegel, R.L., K.D. Miller, and A. Jemal, Cancer statistics, 2020. CA Cancer J Clin, 2020. 70(1): p. 7–30.

2. Mizrahi, J.D., et al., Pancreatic cancer. Lancet, 2020. 395(10242): p. 2008–2020.

3. Hidalgo, M., Pancreatic cancer. N Engl J Med, 2010. 362(17): p. 1605–17.

4. Zhang, L., S. Sanagapalli, and A. Stoita, Challenges in diagnosis of pancreatic cancer. World J Gastroenterol, 2018. 24(19): p. 2047–2060.

5. Bronstein, Y.L., et al., Detection of small pancreatic tumors with multiphasic helical CT. AJR Am J Roentgenol, 2004. 182(3): p. 619–23.

6. Sultana, A., et al., What Is the Best Way to Identify Malignant Transformation Within Pancreatic IPMN: A Systematic Review and Meta-Analyses. Clin Transl Gastroenterol, 2015. 6: p. e130.

7. Wray, C.J., et al., Surgery for pancreatic cancer: recent controversies and current practice. Gastroenterology, 2005. 128(6): p. 1626–41.

8. Lee, E.S. and J.M. Lee, Imaging diagnosis of pancreatic cancer: a state-of-the-art review. World J Gastroenterol, 2014. 20(24): p. 7864–77.

9. van Dongen, G.A., et al., Immuno-PET: a navigator in monoclonal antibody development and applications. Oncologist, 2007. 12(12): p. 1379–89.

10. Delbeke, D. and W.H. Martin, Update of PET and PET/CT for hepatobiliary and pancreatic malignancies. HPB (Oxford), 2005. 7(3): p. 166–79.

11. Lohrmann, C., et al., Retooling a Blood-Based Biomarker: Phase I Assessment of the High-Affinity CA19-9 Antibody HuMab-5B1 for Immuno-PET Imaging of Pancreatic Cancer. Clin Cancer Res, 2019. 25(23): p. 7014–7023.

12. Poruk, K.E., et al., The clinical utility of CA 19-9 in pancreatic adenocarcinoma: diagnostic and prognostic updates. Curr Mol Med, 2013. 13(3): p. 340–51.

13. Schmiegel, W.H., et al., Monoclonal antibody-defined human pancreatic cancer-associated antigens. Cancer Res, 1985. 45(3): p. 1402–7.

14. Takiyama, Y., et al., Reactivity of CO17-1A and B72.3 in benign and malignant pancreatic diseases. Hum Pathol, 1989. 20(9): p. 832–8.

15. Grant, A.G., et al., The generation of monoclonal antibodies against human pancreatic exocrine cancer: a study of six different immunisation regimes. Br J Cancer, 1985. 52(4): p. 543–50.

16. Gao, D. and Y. Chen, Organoid development in cancer genome discovery. Curr Opin Genet Dev, 2015. 30: p. 42–8.

17. Tiriac, H., et al., Organoid Profiling Identifies Common Responders to Chemotherapy in Pancreatic Cancer. Cancer Discov, 2018. 8(9): p. 1112–1129.

18. Somerville, T.D.D., et al., TP63-Mediated Enhancer Reprogramming Drives the Squamous Subtype of Pancreatic Ductal Adenocarcinoma. Cell Rep, 2018. 25(7): p. 1741–1755 e7.

19. Castillo, F.J., et al., Hybridoma stability. Dev Biol Stand, 1994. 83: p. 55–64.

20. Sato, Y., et al., Generation of a monoclonal antibody recognizing the CEACAM glycan structure and inhibiting adhesion using cancer tissue-originated spheroid as an antigen. Sci Rep, 2016. 6: p. 24823.

21. Lee, O.J., et al., CEACAM6 as detected by the AP11 antibody is a marker notable for mucin-producing adenocarcinomas. Virchows Arch, 2015. 466(2): p. 151–9.

22. Burtin, P., et al., Antigenic variants of the nonspecific cross-reacting antigen (NCA). J Immunol, 1986. 137(3): p. 839–45.

23. Suzuki, N., et al., Heterogeneity of circulating carcinoembryonic antigen analyzed by sandwich-enzyme immunoassays with different specificities. Cancer Res, 1987. 47(18): p. 4782–7.

24. Duxbury, M.S., et al., CEACAM6 is a novel biomarker in pancreatic adenocarcinoma and PanIN lesions. Ann Surg, 2005. 241(3): p. 491–6.

25. Perk, L.R., et al., (89)Zr as a PET surrogate radioisotope for scouting biodistribution of the therapeutic radiometals (90)Y and (177)Lu in tumor-bearing nude mice after coupling to the internalizing antibody cetuximab. J Nucl Med, 2005. 46(11): p. 1898–906.

26. Heskamp, S., et al., ImmunoSPECT and immunoPET of IGF-1R expression with the radiolabeled antibody R1507 in a triple-negative breast cancer model. J Nucl Med, 2010. 51(10): p. 1565–72.

27. Boj, S.F., et al., Organoid models of human and mouse ductal pancreatic cancer. Cell, 2015. 160(1-2): p. 324–38.

28. Chakraborty, S. and S. Singh, Surgical resection improves survival in pancreatic cancer patients without vascular invasion-a population based study. Ann Gastroenterol, 2013. 26(4): p. 346–352.

29. Miyabayashi, K., et al., Intraductal transplantation models of human pancreatic ductal adenocarcinoma reveal progressive transition of molecular subtypes. Cancer Discov, 2020.

30. Niu, G., et al., Molecular targeting of CEACAM6 using antibody probes of different sizes. J Control Release, 2012. 161(1): p. 18–24.

31. Strickland, L.A., et al., Preclinical evaluation of carcinoembryonic cell adhesion molecule (CEACAM) 6 as potential therapy target for pancreatic adenocarcinoma. J Pathol, 2009. 218(3): p. 380–90.

32. Moek, K.L., et al., Theranostics Using Antibodies and Antibody-Related Therapeutics. J Nucl Med, 2017. 58(Suppl 2): p. 83S–90S.

33. Castell, F. and G.J. Cook, Quantitative techniques in 18FDG PET scanning in oncology. Br J Cancer, 2008. 98(10): p. 1597–601.

34. Meyer, J.P., et al., Click Chemistry and Radiochemistry: The First 10 Years. Bioconjug Chem, 2016. 27(12): p. 2791–2807.

35. Cherry, S.R., et al., Total-Body PET: Maximizing Sensitivity to Create New Opportunities for Clinical Research and Patient Care. J Nucl Med, 2018. 59(1): p. 3–12.

36. Kloppel, G., F. Petersen, and P.G. Lankisch, Chronic pancreatitis imitating pancreatic carcinoma. Gastrointest Endosc, 1998. 48(2): p. 231.

37. Ruszniewski, P., et al., The diagnostic dilemmas in discrimination between pancreatic carcinoma and chronic pancreatitis. Gut, 2004. 53(5): p. 771.

38. Collisson, E.A., et al., Subtypes of pancreatic ductal adenocarcinoma and their differing responses to therapy. Nat Med, 2011. 17(4): p. 500–3.

39. Almagro, J.C., et al., Progress and Challenges in the Design and Clinical Development of Antibodies for Cancer Therapy. Front Immunol, 2017. 8: p. 1751.

40. Ya, Z., et al., Mouse model for pre-clinical study of human cancer immunotherapy. Curr Protoc Immunol, 2015. 108: p. 20 1 1–20 1 43.

41. Chiou, S.H., et al., Pancreatic cancer modeling using retrograde viral vector delivery and in vivo CRISPR/Cas9-mediated somatic genome editing. Genes Dev, 2015. 29(14): p. 1576–85.

42. Greenfield, E.A., Antibodies : a laboratory manual. 2014.

43. Verel, I., et al., 89Zr immuno-PET: comprehensive procedures for the production of 89Zr-labeled monoclonal antibodies. J Nucl Med, 2003. 44(8): p. 1271–81.

44. Holland, J.P., Y. Sheh, and J.S. Lewis, Standardized methods for the production of high specific-activity zirconium-89. Nucl Med Biol, 2009. 36(7): p. 729–39.

45. Holland, J.P., et al., 89Zr-DFO-J591 for immunoPET of prostate-specific membrane antigen expression in vivo. J Nucl Med, 2010. 51(8): p. 1293–300.

46. Ross, P.L., et al., Multiplexed protein quantitation in Saccharomyces cerevisiae using amine-reactive isobaric tagging reagents. Mol Cell Proteomics, 2004. 3(12): p. 1154–69.

47. Perkins, D.N., et al., Probability-based protein identification by searching sequence databases using mass spectrometry data. Electrophoresis, 1999. 20(18): p. 3551–67.

