## Supplementary Figures for "Detection of incipient pancreatic cancer with novel tumor-specific antibodies in mouse models"

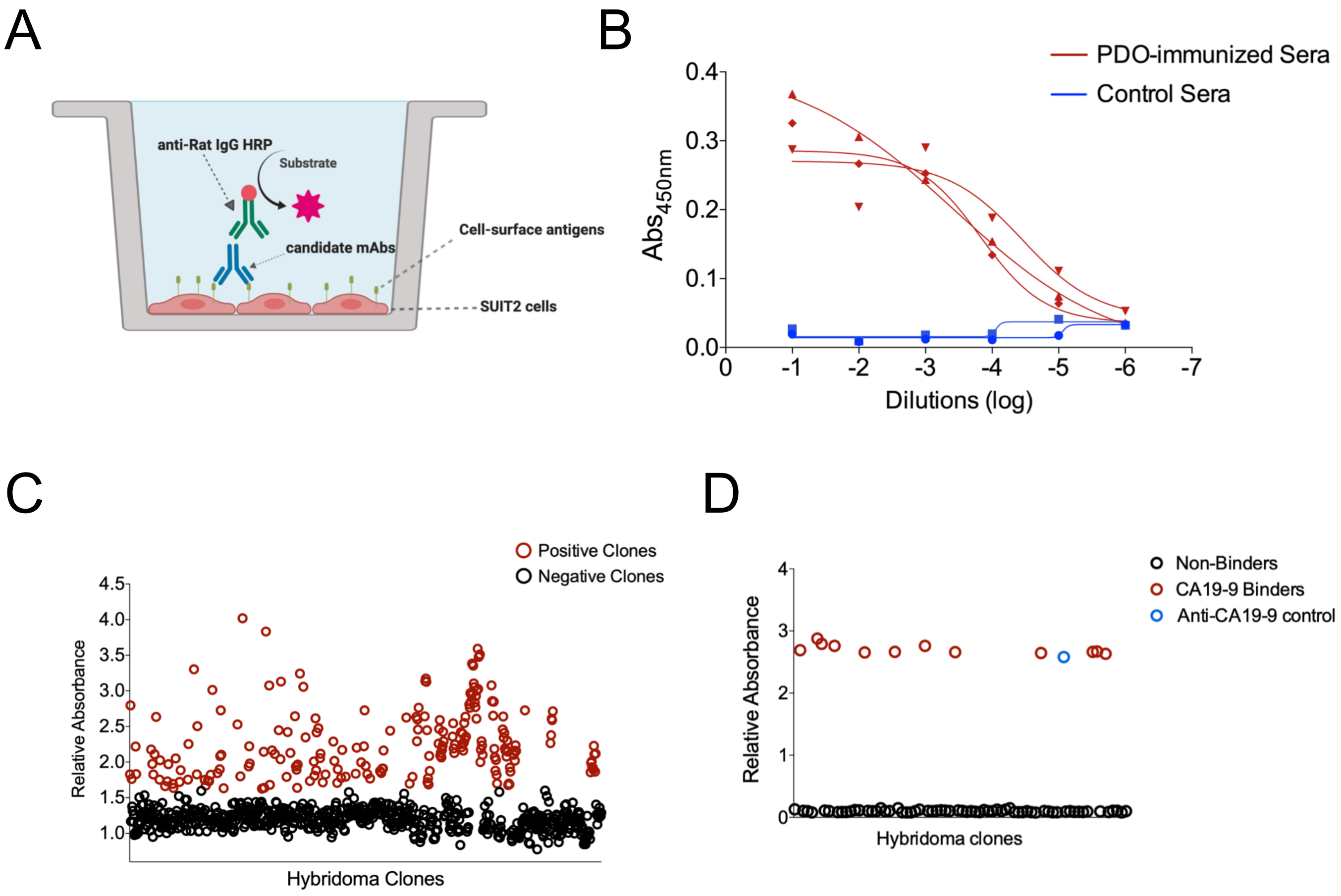


Supplementary Figure 1

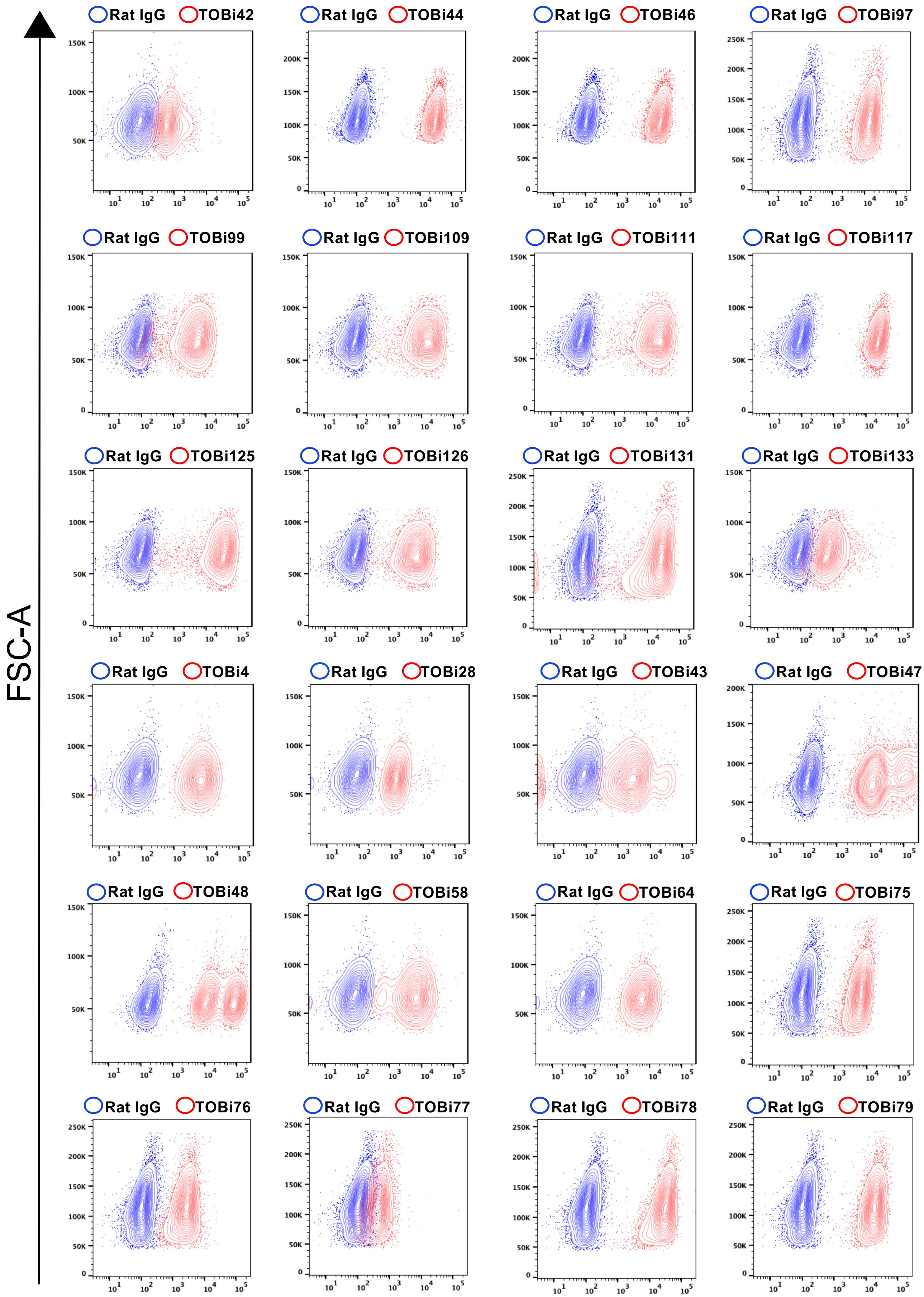


Supplementary Figure 2

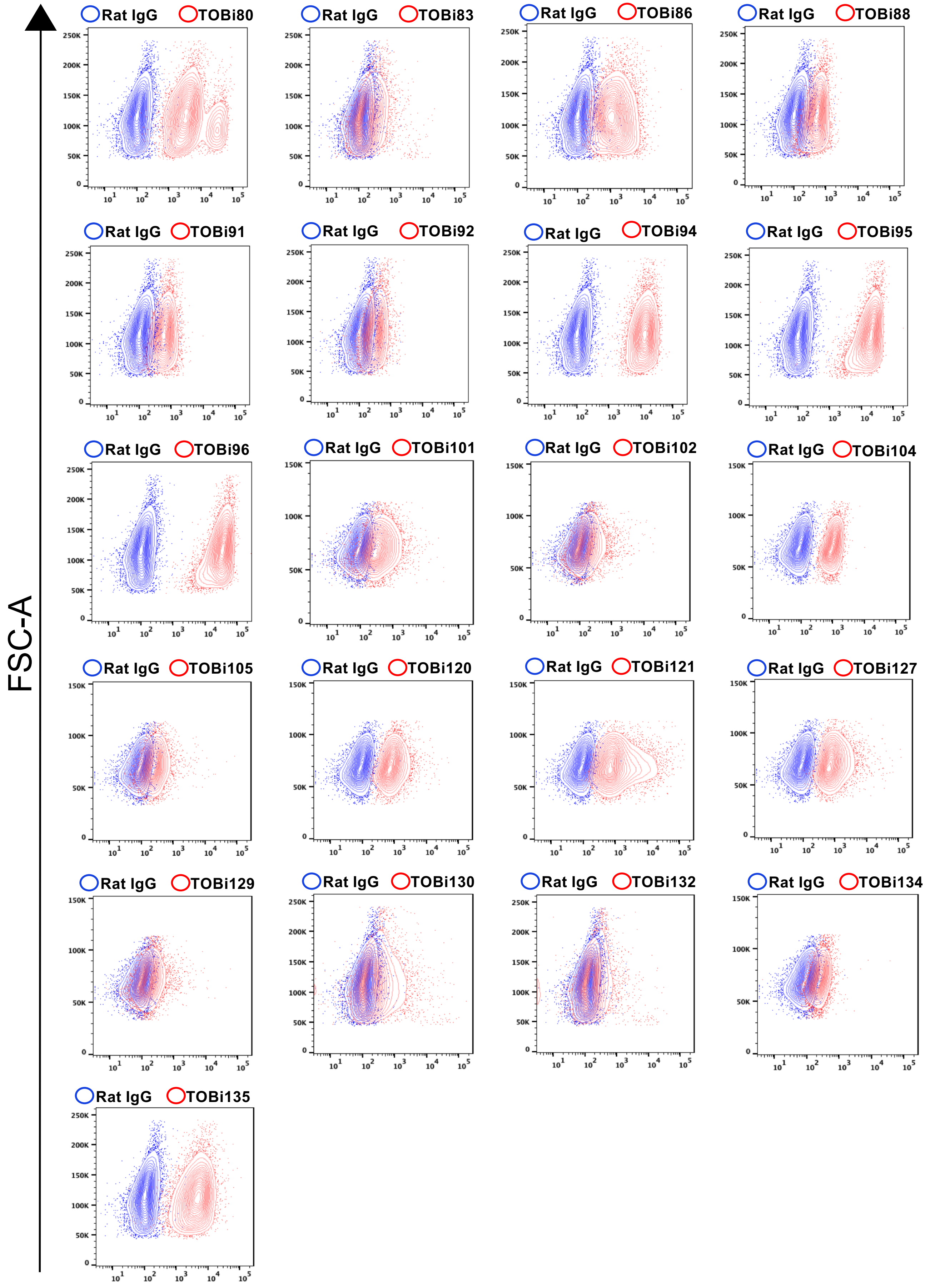


Supplementary Figure 3

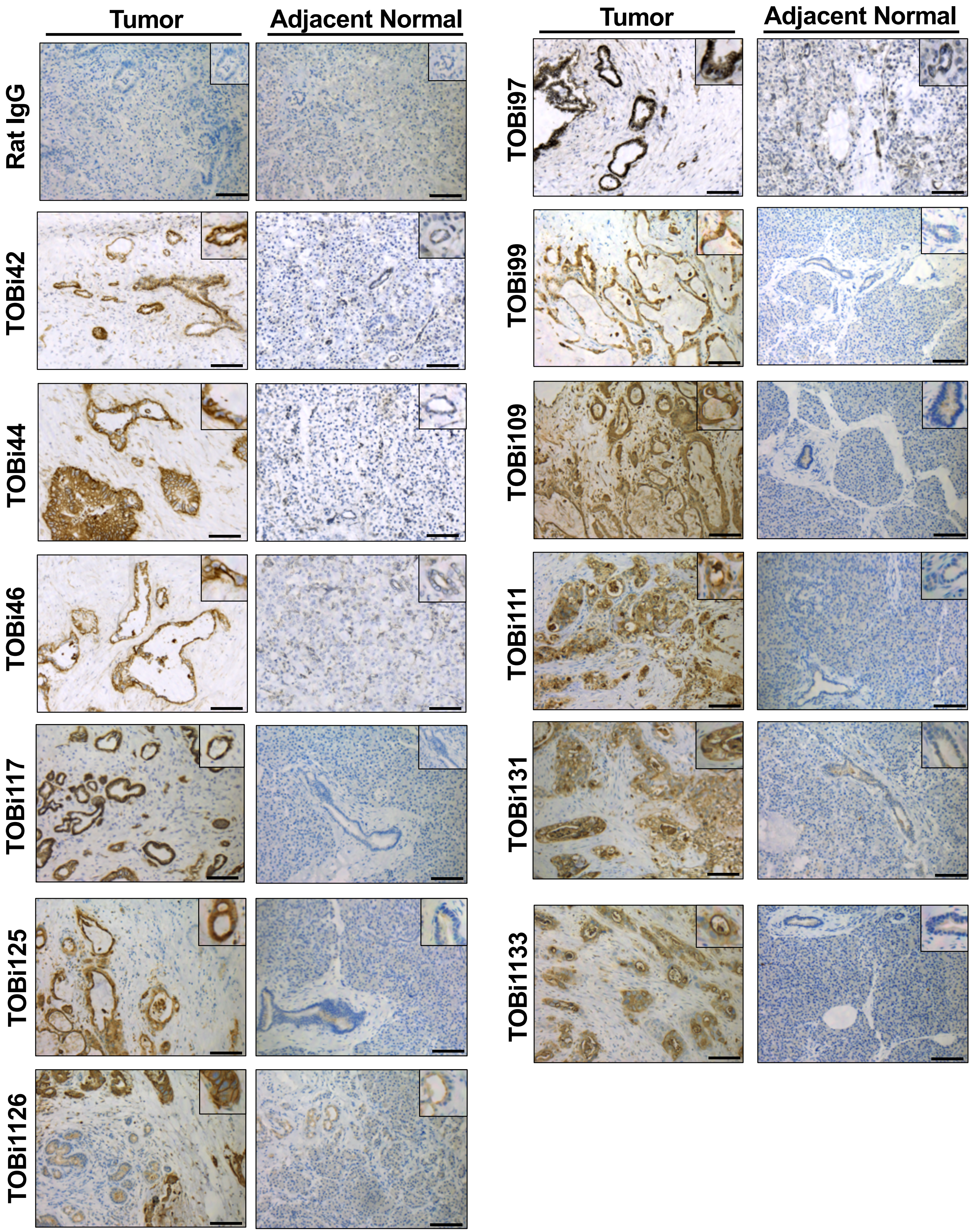


Supplementary Figure 4

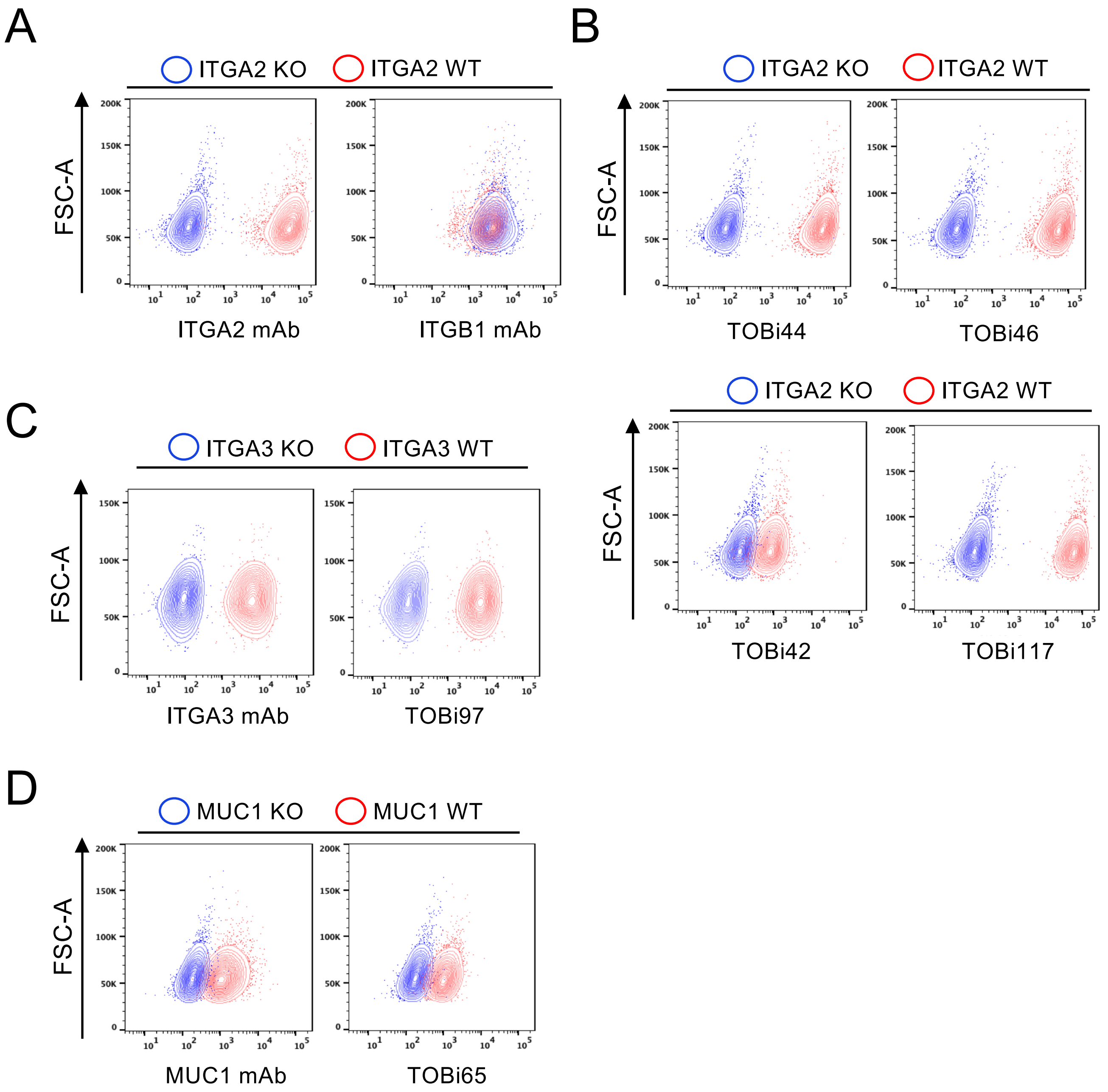


Supplementary Figure 5

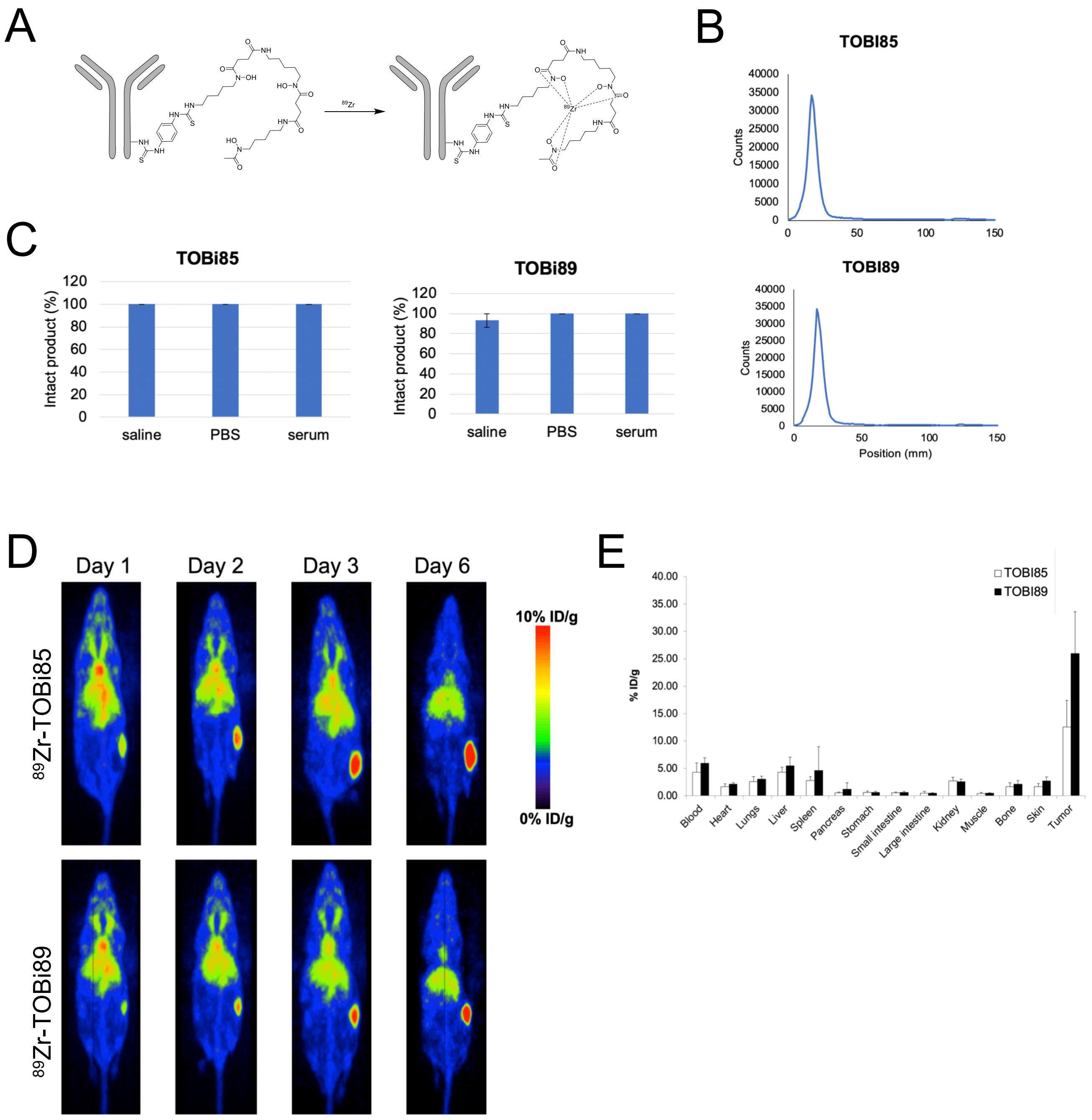


Supplementary Figure 6

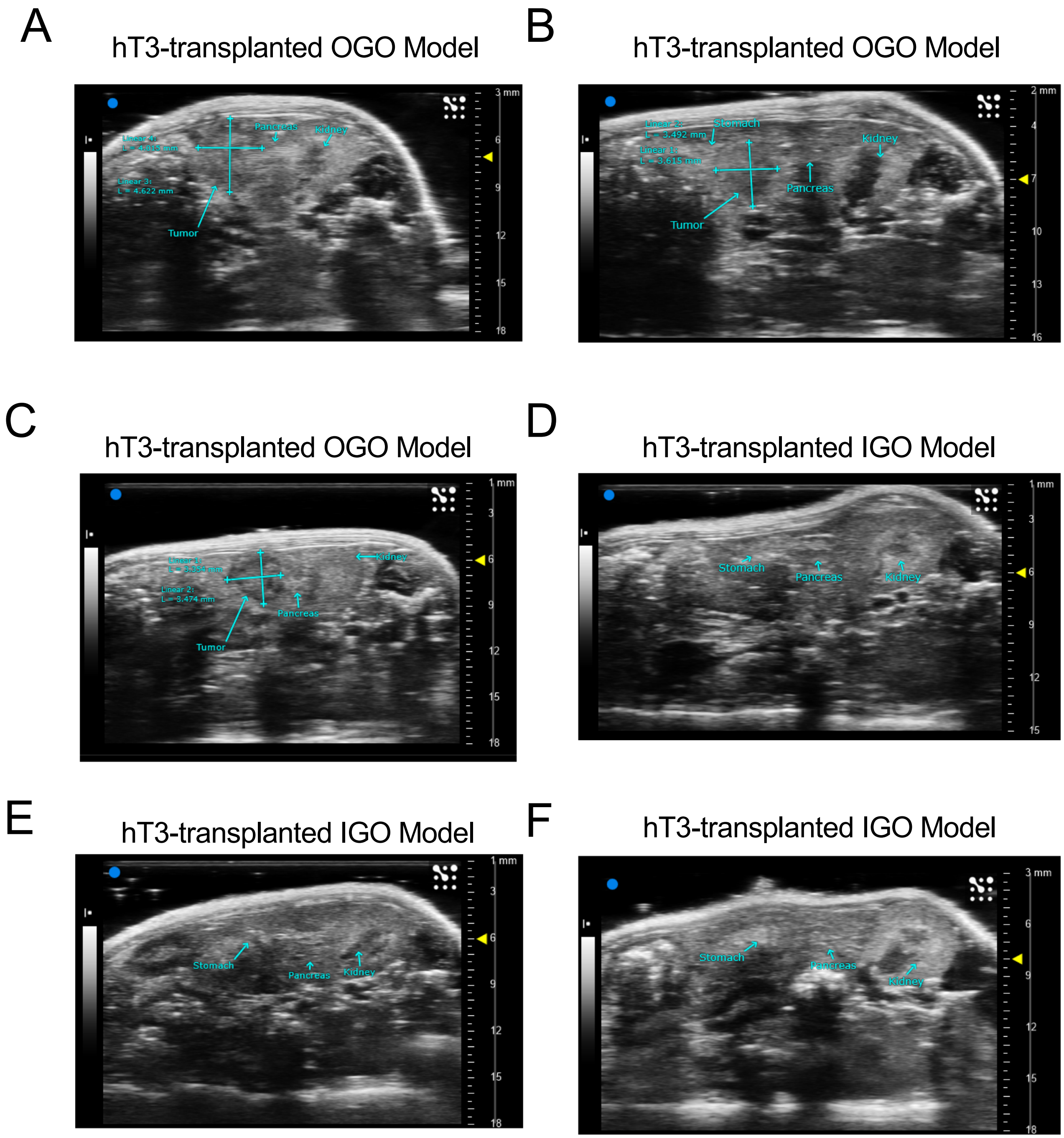


Supplementary Figure 7

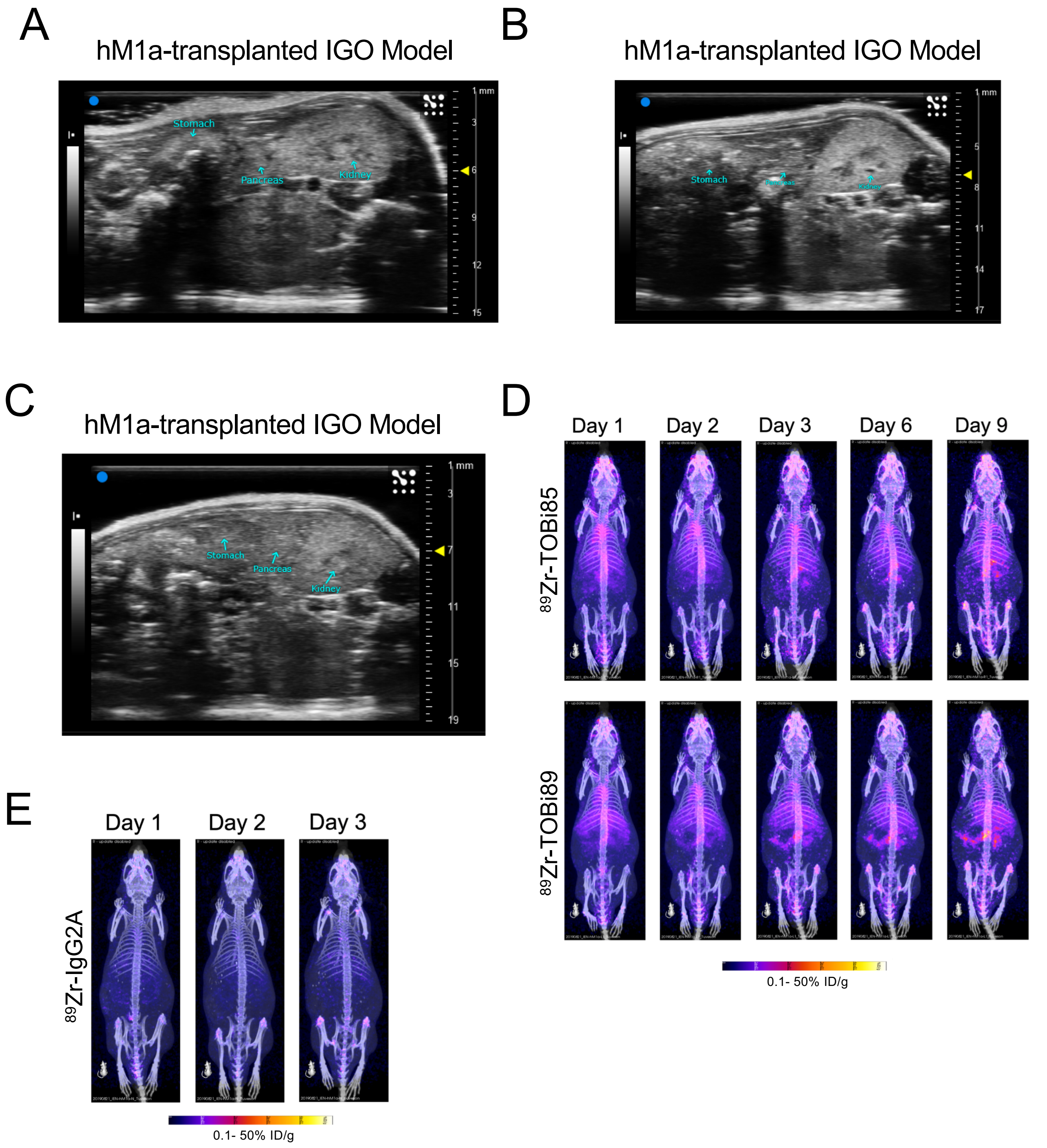


Supplementary Figure 8
